# Cuticular wax, but not the cutin matrix, is renewed during the lifespan of *Clusia rosea* leaves *^13^CO_2_ labelling and gas exchange study*

**DOI:** 10.1101/2022.09.26.509460

**Authors:** Jiří Kubásek, Tereza Kalistová, Jitka Janová, Balzhan Askanbayeva, Jan Bednář, Jiří Šantrůček

## Abstract

- The plant cuticle, which aided the water-to-land transition of plants, provides various services to the plant surface, and its synthesis and maintenance represent substantial metabolic costs. Nevertheless, only limited information regarding cuticle dynamics is available.
- We determined the composition and dynamics of *Clusia rosea* cuticular waxes and matrix using ^13^CO_2_ labelling, compound-specific and bulk isotope ratio mass spectrometry. Collodion was used for wax collection; gas exchange techniques were employed to test for any collodion effects on living leaves.
- Cutin matrix (MX) area density did not vary between young and mature leaves and between leaf sides. Only young leaves incorporated new carbon into their MX. Collodion-based sampling discriminated between epicuticular (EW) and intracuticular wax (IW) effectively. EW differed in composition from IW. The newly synthetized wax was deposited in IW first and later in EW. Both young and mature leaves synthetized IW and EW; the faster dynamics in young leaves was not due to a faster synthesis rate but was the result of lower wax coverage. Longer-chain alkanes were deposited preferentially on the abaxial, stomatous leaf side, producing differences between leaf sides in wax composition.
- We introduce a new, sensitive isotope labelling method and demonstrate that cuticular wax is renewed during leaf ontogeny of *Clusia rosea*. We discuss the ecophysiological significance of the new insights.

## Introduction

The plant cuticle is a multifunctional, hydrophobic biofilm usually studied on aboveground plant surfaces (Riederer & Muller, 2006) but found recently also on the root cap (Berhin *et al*., 2019). With an overall thickness of only about 1 μm in mesophytic leaves (Jeffree, 2018), its excellent gas barrier properties and self-cleaning ability have fascinated plant scientists and bioengineers for many decades (Skoss, 1955; Barthlott & Neinhuis, 1997; Fernández *et al*., 2017). Modern cuticles serve not only as a barrier but also as a communication interface between plants and their environment, as well as a developmental regulator; however, cuticles very probably arose primarily to aid plants’ transition from water to land, by limiting uncontrolled water loss, about 450 million years ago (Kerstiens, 1996b; Niklas *et al*., 2017; Kong *et al*., 2020). In concert with gas-filled intercellular spaces (ICSs) and adjustable gas valves – stomata – the cuticle enabled homoiohydry (i.e. decoupling of plant water regime from the external water potential) in plant sporophytes spanning from moss capsules to the biggest trees. Homoiohydry was hypothesized to be a prerequisite of large plant size and high productivity (Proctor & Tuba, 2002) and thus to support the rich metazoan life including humanity.

A thin (rudimental) cuticle was found on many bryophyte gametophytes (where no stomata and ICSs are present) without any clear function (Matos *et al*., 2021). On the other hand, moss sporophytes (Budke & Goffinet, 2016; Kubásek *et al*., 2021) and vascular plant leaves and stems as well as various flower organs and fruits bear well-formed cuticles with very low conductance to water and CO_2_ diffusion (Woolley, 1967; Boyer, 2015). Many studies measured environmental dependencies of cuticle properties (Baker, 1974; Kerstiens, 1996a, 1997). Increased relative air humidity (Schönherr & Schmidt, 1979; Schreiber *et al*., 2001) and temperature (Riederer & Schreiber, 2001) have been found to increase cuticle conductance, but no correlation between this conductance and cuticle thickness, structure or composition has been found in large sets of plant species (Kerstiens, 1996c; Hauke & Schreiber, 1998; Riederer & Schreiber, 2001; Zeisler-Diehl *et al*., 2017).

Even though the plant cuticle was studied as early as 1830 by Brongniart (Skoss, 1955), much remains still unknown. The fine structure of the plant cuticle is complex and variable (Jeffree, 2018). A simplified view distinguishes a polyester cutin matrix (MX), which consists of covalently bound C16, C18 hydroxy-fatty acids (and their derivatives), glycerol and phenolics, forming a 3D scaffold (Fich *et al*., 2016). The MX is impregnated with a variety of aliphatic and cyclic lipids (intracuticular wax, IW) and covered by a purely lipidic layer on the uppermost cuticle surface (epicuticular wax, EW; Haas & Rentschler, 1984; Jetter & Schäffer, 2001a). Many plant species also incorporate cutan (non-ester bound moieties) into their MX, and almost all species studied so far also accommodate various non-lipidic molecules (e.g. flavanols, phenolics) in small to dominant proportions (Karabourniotis *et al*., 2001; Jetter *et al*., 2018; Kong *et al*., 2020). Waxes may be largely extracted using organic solvents (e.g. chloroform) and their removal leads to an increase in cuticular permeability for water by two or three orders of magnitude (Schönherr & Schmidt, 1979). The ultrastructure of the cuticle is even more complex. Cuticle proper, the outermost cutin-rich layer, descending from the embryonal procuticle, and cuticle layer, the polysaccharide-rich fraction in contact with the epidermal cell wall, are well recognizable via transmission electron microscopy (TEM) in many plant species. Moreover, electron-lucent vs. electron-dense micro-domains may be found in each layer (Jetter *et al*., 2000; Jeffree, 2018). Traditionally, the cuticle is considered as a distinct layer on the epidermis since it may be isolated enzymatically (Skoss, 1955; Vráblová *et al*., 2020). A more recent view, however, perceives a continuum of cell wall and cuticle (Guzmán *et al*., 2014; Fernández *et al*., 2016).

We currently have nearly complete pictures of the biosynthesis of many cuticle compounds (cutin monomers, triterpenes, phenolics), although restricted to several model species, e.g. *Arabidopsis* (Wen & Jetter, 2009; Bernard & Joubès, 2013; Wang *et al*., 2020), tomato (Yeats *et al*., 2012) or maize (Petit *et al*., 2017). Much less is known, for example, about cutin monomer export and polymerization (Fich *et al*., 2016; Niklas *et al*., 2017), lipid export to the apoplast (Pighin *et al*., 2004; Panikashvili *et al*., 2011) and assembly of the functional cuticle (Samuels *et al*., 2008; Bourgault *et al*., 2020; Seale, 2020).

Probably the scarcest domain in our current understanding is cuticle dynamics. Gas chromatography – mass spectrometry revealed that the total wax amount and its composition change during development of leaves or other organs (Hauke & Schreiber, 1998; Jetter *et al*., 2018) and that IW and EW may be chemically distinct (Jetter & Schäffer, 2001a). Changes in environmental factors as well as internal plant signaling are known to affect the cuticle quickly (Skoss, 1955; Baker, 1974; Koch *et al*., 2004). For instance, just seven day of irrigating *Lepidium sativum* seedlings with 10^−4^ M abscisic acid (ABA) led to a shift in cuticular wax to compounds with longer chains (Macková *et al*., 2013). Atomic force microscopy and scanning electron microscopy have also proved useful for the study of EW regeneration (Neinhuis *et al*., 2001; Koch *et al*., 2004), revealing distinct groups of plants that differ in their ability to regenerate their EW (Neinhuis *et al*., 2001).

Surprisingly, stable isotope labelling techniques have only been used in a few cuticular studies so far. Mostly, a heavy water (D_2_O) irrigation approach was applied with the aim to detect whether and how fast cuticle components are renewed (Kahmen *et al*., 2011; Gao *et al*., 2012). However, the results remain controversial. It is not clear if wax is deposited during the whole leaf lifespan, as shown by (Sachse *et al*., 2009; Gao *et al*., 2012), or only early in leaf ontogeny (Flore & Bukovac, 1974; Kahmen *et al*., 2011). Compound-specific analyses revealed that turnover of cuticular fatty acids (C16 and C18) is much faster (2 to 3 days) than the turnover of long-chain lipids (5 to 128 days; (Gao *et al*., 2012). This may be the result of a non-cuticular origin of those fatty acids (see discussion). The dynamics of cutin MX as well as various wax compounds remain unknown.

The abovementioned gaps are often related to the methods applied. Whole-leaf immersion in solvents (Medina *et al*., 2004) and/or extraction of total leaf lipids may yield biased results with respect to the genuine lipid spectra of the cuticle. The GC-MS separation techniques may distinguish cuticular compounds thereafter, but their origin – IW, EW or lipids still present in ER of epidermal cells – cannot be distinguished. Thus, here we sampled EW and IW separately, using selective techniques.

*Clusiaceae* is a tropical family of some 37 genera and 1610 species (very close to *Hypericaceae*) in the big order Malpighiales (Gustafsson *et al*., 2002). Physiological and cuticular studies exist for many species in the genus *Clusia* (Medina *et al*., 2004; Medina *et al*. 2006; Lüttge, 2008). There is a tendency for environmentally-induced CAM metabolism in most *Clusia* species (Lüttge, 2008). *Clusia rosea* Jacq. is nowadays frequently cultivated in glasshouses and grown in frost-free outdoor sites worldwide. We chose this species for five reasons: 1/ it grows vigorously and CO_2_ assimilation is substantial under optimal conditions; 2/ leaves are large, relatively thick and EW may be sampled easily without damage to the leaf; 3/ leaves are hypostomatous and so, stomatous abaxial and astomatous adaxial cuticles may be compared; 4/ wax composition is well defined with several compounds prevailing in the total wax (Medina *et al*., 2004; Medina *et al*., 2006); 5/ the cuticles may be isolated enzymatically and IW extracted subsequently.

Our aim was to reveal age-dependent and side-specific dynamics of the main cuticular wax compounds and the cutin matrix. We identified and quantified dominant EW and IW compounds in young and mature leaves of *Clusia rosea*, separately for adaxial and abaxial cuticles. We used the ^13^CO_2_ pulse-chase approach to follow dynamics of deposition of the major wax compounds in EW, IW and cutin matrix. Methodologically, we tested the ability of collodion solution to discriminate between EW and IW. The impact of collodion treatment on leaf gas exchange and stomatal behavior was also evaluated.

## Material and Methods

### Plant material

Commercially obtained plants of autograph tree (*Clusia rosea* Jack.) were grown in a greenhouse (University of South Bohemia, České Budějovice, 48.9779119N, 14.4457319E) for two years before the experiments. Temperature was maintained between 20 and 26 °C during the day and at least 18 °C at night. Relative humidity was 50 to 80 %. The natural photoperiod at the time of the experiment (February to March 2020, ≈ 12 h) was extended to approximately 16 h using metal halide high-pressure lamps with PPFD at ≈ 150 μmol photons m-_2_ s^−1^.

### ^13^CO_2_ Labelling and sampling strategy

Four plants (about 30 cm tall) were transferred to a gas-tight Plexiglas box (internal dimensions were 60 × 60 × 60 cm and volume 190 L) for 2 h of assimilation in a ^13^CO_2_ -enriched atmosphere (Fig. S1). A fan inside the box reduced the resistance of the boundary layer. The box was placed in a cultivating room where the temperature was maintained at 20 °C. PPFD irradiation of ≈400 μmol photons m-_2_ s^−1^ was supplied by linear LED sources, two units above and two more on each side of the box, to achieve the most homogeneous light possible (see Fig. S2 for spectral output). At the beginning, we let the open box equilibrate with the atmosphere of the room. After a few minutes, steady state was reached, the box was hermetically sealed, and about 95 ml of ^13^CO_2_ (>99% atoms, Sigma-Aldrich, Luis., USA) was injected through a septum in the box wall, yielding ≈800 μmol CO_2_ mol^−1^ (twice the ambient level and about half of which is ^13^CO_2_). The plants assimilated CO_2_ for two hours. Every 30 min, the box was opened, a new equilibrium was reached, and new ^13^CO_2_ applied after closure. We previously found that the same plants were able to reduce the CO_2_ concentration inside by ≈30 % within 30 min (using a 6400XT unit, LiCor, Lincoln, NE, USA, and isotopically normal CO_2_). After 2 h, the plants were transferred back to the greenhouse for the remainder of the experiment.

The overall sampling design was as follows: One young, more or less fully expanded leaf from the current year and one mature leaf from the previous year (typical projected leaf area 30 to 60 cm_2_) were collected from each plant before labelling (t=0), immediately thereafter (t=2 h), and 12, 24, 48, 96, 168 (one week), 336 (two weeks), and 504 h (three weeks) later. Epicuticular wax (EW) was removed separately from the adaxial and abaxial surfaces immediately after leaf excision (see below). One 2 cm diameter disc (3.14 cm^2^) was then cut and dried at 80 °C for 12 h to obtain dry mass, and two to four discs of the same size (depending on leaf area) were cut to isolate the cuticle. All this work was done within one hour of harvest. Finally, the rest of the leaves were placed in a freezer (-80 °C) for subsequent analysis of water-soluble compounds as a putative source for lipid biosynthesis.

### Isolation of epicuticular and intracuticular waxes and wax-free cuticular matrices

EW was removed from each side of the leaf using a collodion solution (Haas & Rentschler, 1984) applied with a fine brush in two successive applications. The strips were peeled off after polymerization (≈1 min) and extracted in 4 mL of n-hexane in 20-mL scintillation vials on a roller overnight at room temperature (n-hexane was found to have comparable efficiency in extracting various compounds from collodion but not from cuticle, where chloroform was more efficient). It was then concentrated by evaporation under a gentle stream of air to 1 ml and transferred to 2 ml chromatographic vials for GC-IRMS analyses. The two strips from each side were able to remove approximately 90 % of the EW in the pilot experiment (Fig. S3), and we therefore chose this protocol as a good compromise between efficiency and leaf damage. Two to four discs 2 cm in diameter were punched from EW-free leaves, and the cuticle was isolated enzymatically in a 2 % mixture of cellulase and pectinase (Schönherr & Riederer, 1986). Isolated cuticles were separated into adaxial and abaxial ones (using a microscope) and IW was extracted in 1 ml chloroform in 2 ml vials at room temperature on a roller overnight. Then, the wax-free cuticular matrices (MX) were removed, dried, weighed (using a Mettler Toledo MT5 microbalance, Columbus, OH, USA) and analyzed for ^13^C content using an elemental analyzer (Flash 2000, Thermo, Bremen, Germany) coupled to isotope MS (Delta XL, Thermo, Bremen, Germany).

### Gas chromatography, mass spectrometry and stable isotope analyses

The EW and IW extracts obtained with collodion were analyzed on a gas chromatograph (Trace 1310, Thermo, Bremen, Germany). A Restek Rxi-5MS-Sil column (30 m x 0.25 mm x 0.25 μm film thickness) was used and the flow rate was 1.5 ml min^−1^ of helium as carrier gas. The injection (at 300 °C) was splitless for 1.5 min, then split flow at 100 ml min^−1^ for another 1 min and 5 ml min^−1^ for the rest of the time (gas saver). The oven temperature program was set to 50 °C during injection and for the following 2 min, then increased with a gradient of 40 °C min^−1^ to 200 °C, then at 4 °C min^−1^ to 310 °C, and was isothermal at 310 °C for the rest of the analysis (c. 55 min in total). To maintain isotopic fidelity and avoid introducing additional isotopic noise, we chose to measure non-derivatized samples. The eluting compounds were oxidized to CO_2_ via IsoLink II interphase (Thermo, Bremen, Germany) at 1000 °C and introduced into a continuous flow isotope ratio MS (Delta V Advantage, Thermo, Bremen, Germany). The internal standard n-tetracosane (C24 alkane) calibrated at ^13^C against VPDB was added to the samples at a concentration of 10 or 20 μg ml^−1^ to quantify the compounds and check for isotope shifts and/or potential memory effects. In addition, a GC calibration curve was developed for C10 to C40 alkanes to correct for the decrease in GC sensitivity in high-boiling compounds. Finally, isotopic consistency across different carbon chain lengths was monitored using Arndt Schimmelmann’s A7 isotope mixture of C16 to C30 n-alkanes (Schimmelmann *et al*., 2016). For compound identification, the same setup and a capillary column with a GC-APCI-MS system (456-GC + Compact QToF; Bruker Daltonics; Germany) were used. Compounds were identified using available standards and/or accurate molecular fragment masses. We also measured a subset of derivatized samples of both EW and IW from young and mature leaves (50 μL BSTFA + 100 μL pyridine, 2 h at 80 °C; Hauke & Schreiber, 1998). EW did not contain additional compounds in substantial amounts (< 1 % of total peak area) whereas IW of young leaves revealed some primary alcohols.

### Isolation and measurement of water-soluble leaf carbon

To obtain the ^13^C content of the metabolically active pool, we isolated the water-soluble organic fraction of the leaves. Leaves stored in a deep freezer (-80°C) were thawed, dried, and homogenized to a fine powder using a ball mill (MM400, Retsch, Haan, Germany). Approximately 100 mg of this powder was mixed with 1 ml of deionized water, sonicated at room temperature for 2 h at 80 °C and finally centrifuged at 10,000 g for 10 min. The supernatant was gently dropped into tin capsules and left to evaporate until the required dry weight (≈ 100 μg) was obtained. Samples were analyzed for ^13^C content by combustion using an elemental analyzer (Flash 2000, Thermo, Bremen, Germany) coupled to isotope ratio MS (Delta XL, Thermo, Bremen, Germany).

### Gas exchange measurements before and after collodion treatment

Water and CO_2_ exchange was measured separately for the adaxial and abaxial side of the leaf using two Li-6400XT units (LiCor, Lincoln, NE, USA) arranged “in tandem” (see Fig. S4 for more details on the setup of the two Li-6400XT units). Ambient conditions in the leaf chamber were as follows: 25 °C, 400 μmol mol^−1^ CO_2_, and 40 % to 60 % relative humidity. A leaf acclimated in the dark for 1 h was placed in the leaf chamber. After a few minutes in the dark (to reach the steady state of the system), the light was switched to 1000 μmol m-_2_ s^−1^, and CO _2_ assimilation and stomatal conductance were recorded each 10 s for one hour, separately for the adaxial and abaxial side. The leaf was then removed from the chamber and both sides treated with collodion as mentioned above. Gas exchange measurements were repeated on the same leaf approximately 24 h later (Fig. S5).

### Calculations and statistics

^13^C excess (At%) of any particular labelled compound or structure is defined as the difference between the ^13^C content after and before labelling. The variability in the natural content of ^13^C in lipid substances (1.063 ± 0.003 At%) was small in comparison with the labelling-induced range (up to 1.5 At%), which ensured high sensitivity of the method.

We measured ^13^C excess in water-soluble leaf carbon (SolC) as a proxy of the metabolic pool used in the synthesis of cutin and wax precursors (SolC values are able to even out differences between leaves in their labelling intensity over time).

Then, new wax deposition (NWD, μg cm-_2_) may be calculated as:

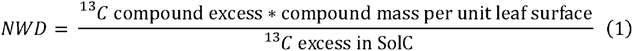

Statistical significance was tested using one-way or two-way factorial ANOVA followed by Tukey’s post-hoc HSD test. When normality and/or homoskedasticity (Shapiro-Wilks test) was violated (typically for EW wax load), a logarithmic transformation was used and log-transformed data tested instead.

## Results

### Leaf and cuticle characteristics

Leaf area was about 35 % smaller and stomatal density 27 % greater in young leaves than in mature leaves, indicating an ongoing slight expansion of young leaves. EW formed ‘floes’ on the leaf surface visible by cryo-SEM (Fig. 1). The amount of collodion-collected EW varied considerably and had a strongly and positively skewed distribution (Table 1, Fig. S6). Young leaves had a much lower EW load (0.20-0.41-1.58 μg cm-^2^ [q25%-median-q75%, N=144]) than mature leaves (3.1-7.4-27.7 μg cm-^2^). Thus, mature leaves accumulate at least one order of magnitude more EW. Cuticles without EW had an area density of 329 ± 110 μg cm-^2^, which may represent a thickness of about 3.0 μm, assuming a mass density of 1.1 g cm^−3^ (Tsubaki *et al*., 2012; Onoda *et al*., 2012). Statistical distribution fitted well to Gaussian, and neither leaf age nor side had a significant effect on MX area density (ANOVA, F=(1,132)=0.518, p=0.472, N=134). The IW extracted from a subset of isolated cuticles accounted for about 16.5 % and 20 % of cuticular mass in young and mature cuticles, respectively (ANOVA, F=(1,12)=7.67, p=0.017, N=16, Table 1). The full GC data set yielded similar results. Young leaves had a lower IW load (2.1-3.0-4.6 μg cm-^2^, N=144) than mature leaves (3.1-4.9-7.1 μg cm-^2^) and distribution of both data sets were slightly positively-skewed (Fig S6).

**Table 1.**
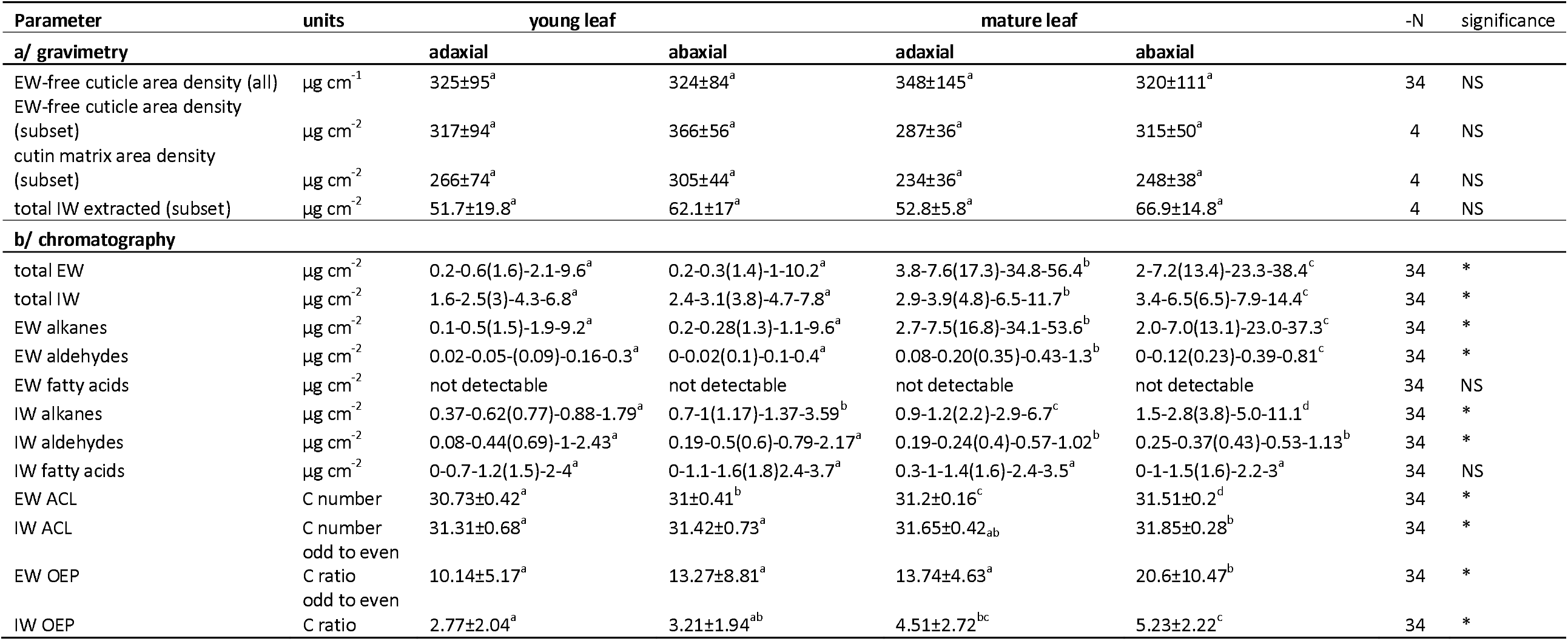
Cuticle and wax characteristics of *Clusia rosea* leaves. Values with normal distributions are shown as mean ± one standard deviation. Non-Gaussian distributions (predominantly positively skewed) are presented as q25%-median(mean)-q75%-q95%. EW and IW denote epicuticular and intracuticular waxes, respectively. Average carbon length, ACL = S([C_i_] x i)/ S[C_i_], where [C_i_] is the concentration of n-alkane containing i carbon atoms, OEP (odd to even preference index) = S[C_i_]odd/ S[C_i_]even, where [C_i_]odd are concentrations of odd carbon numbered compounds from C25 to C37 and [C_i_]even are those of even numbered compounds from C26 to C38. Subset of cuticles is shown where MX and IW load were determined not only chromatographically but also gravimetrically. Subsets with different upper-key letters differ significantly. Significance criteria NS: P > 0.05, * P < 0.05.

**Figure 1.**
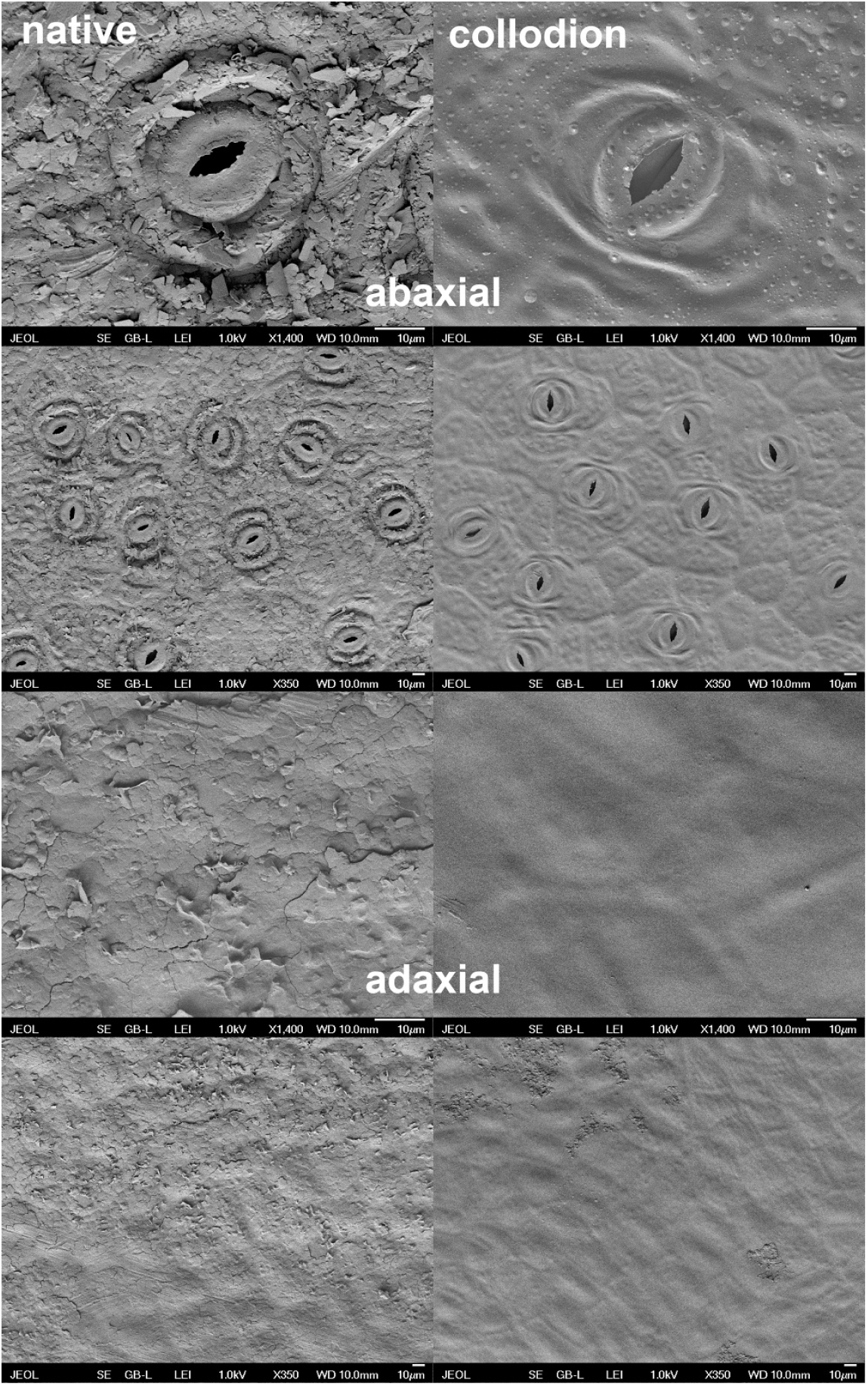
Cryo-SEM images of leaf surfaces of *Clusia rosea* with and without collodion treatment. Intact surfaces (left column) and surfaces after stripping with two layers of collodion applied subsequently (right column). The upper two rows show the abaxial (stomatal), the bottom two rows the adaxial (astomatous) leaf side. Two magnifications are shown (see scale bars).

### Relative abundance of epicuticular and intracuticular wax components

N-alkanes, aldehydes and fatty acids were detected in native (non-derivatized) GC samples (Fig. 2, Fig. S7). C29, C31 and C33 alkanes (n-nonacosane, n-hentriacontane and n-tritriacontane) were predominant in EW and together accounted for 88.8-90.9-93.8 % (q25%-median-q75%) of all GC-amenable compounds, whereas in IW they accounted for 24.6-36.4-50.5 %. The proportion of n-alkanes was always higher in mature cuticles (Table 1, Fig. 2). Aldehydes (C28 to C36) were most abundant in the IW of young leaves (21 %) and rarest in the EW of mature leaves (1.9 %). Fatty acids (C16 and C18) were undetectable in EW but formed a surprisingly high percentage of IW from enzymatically isolated cuticles. Primary alcohols (with an even number of carbons) were also detected in small amounts (mainly in young leaves) in a subset of derivatized samples (not shown).

**Figure 2.**
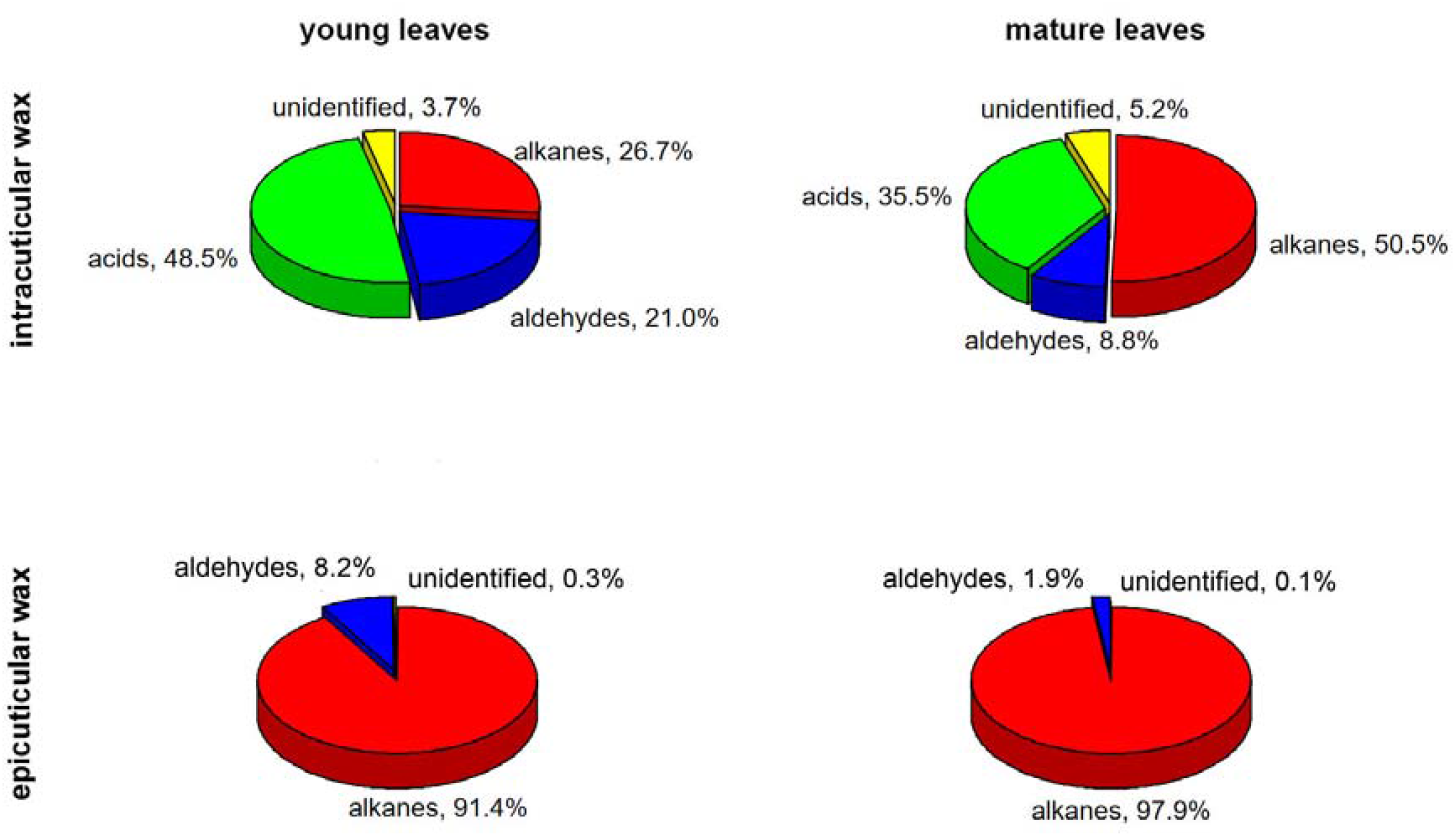
Relative abundance of epicuticular and intracuticular wax compounds of *Clusia rosea* leaves. Percentages of alkanes, aldehydes and fatty acids measured in intracuticular and epicuticular waxes from young and mature leaves are shown. Adaxial and abaxial leaf sides are pooled due to insubstantial differences in overall compound class abundances. See Table 1 for side-specific absolute quantification of each compound class.

Our approach was to detect the dynamics of major compounds. The difficulty of isotopic precision for derivatized and/or minor compounds led us to ignore them in further analyses. The absolute quantification of compound classes can be found in Table 1. Differences between opposite sides of leaves were minor and will be described separately.

### Efficiency of wax sampling by collodion; intracuticular and epicuticular wax dynamics

Cryo-SEM images indicated that two strips of collodion removed the visible wax on the leaf surface (Fig. 1). We tested the effectiveness of collodion for removing EW and its ability to distinguish between EW and IW. The efficiency of three subsequent collodion strips in removing EW can be seen in Fig. S3. Two strips removed at least 90 % of EW (slightly more for alkanes than for semi-polar compounds). We provide two lines of evidence that EW and IW are distinct and may be sampled selectively by collodion. First, individual compounds and compound classes (i.e. alkanes vs. aldehydes) differed in their abundances between EW and IW (Table 1, Fig. 2). Second, assimilated carbon (^13^C) was visible as early as 2 h after labelling in IW with a much later appearance in EW, which became progressively more strongly labelled with time and could surpass IW by the end of the experiment (Fig. 3, Fig. 4).

**Figure 3.**
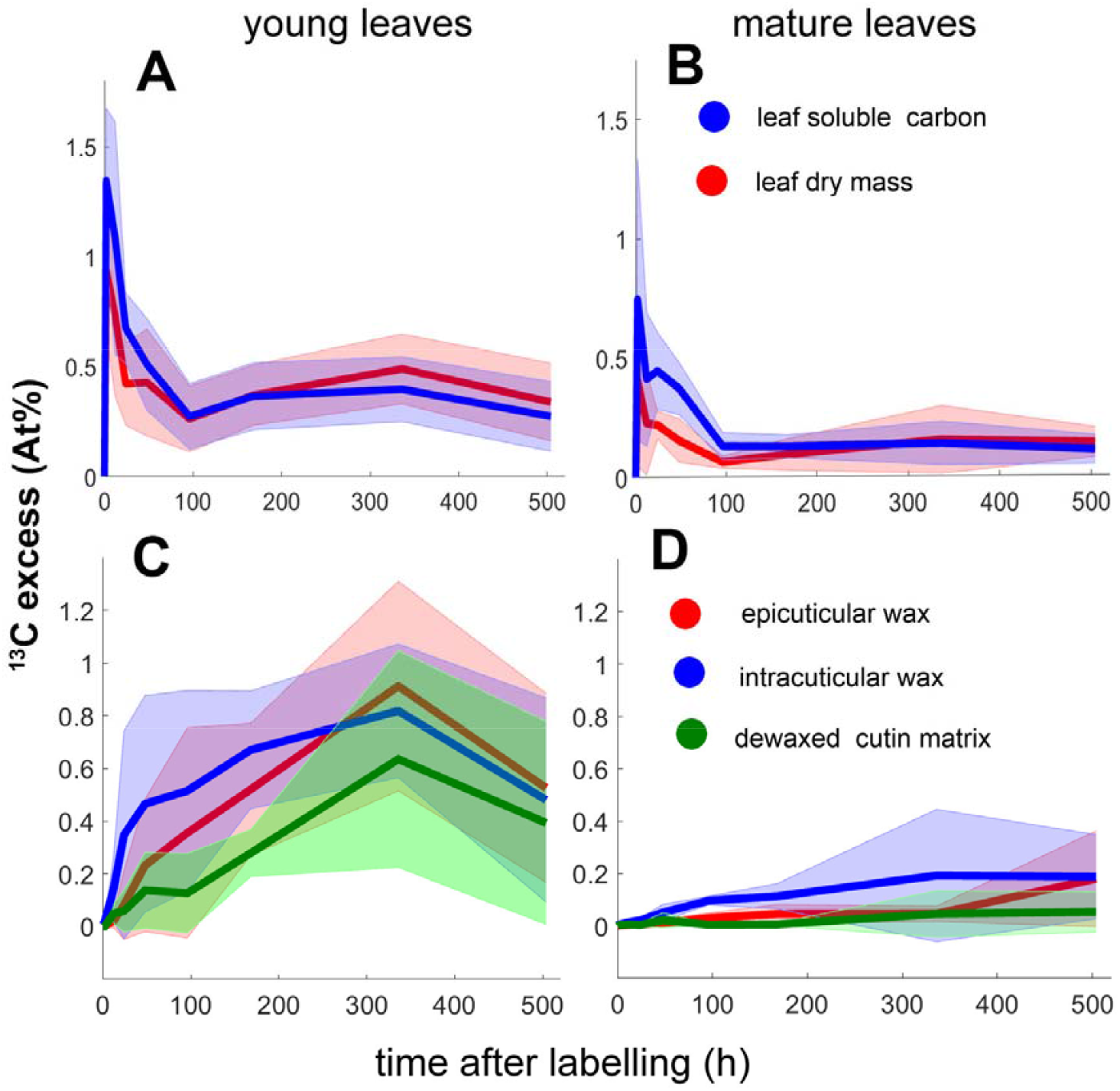
Dynamics of ^13^C abundance in leaf water-soluble fraction, dry mass (A,B) and cuticular constituents (C,D) of *Clusia rosea*. The dynamics is shown for young (A, C) and mature (B, D) leaves before labelling, after 2 h of CO_2_ assimilation in air highly enriched in ^13^CO_2_ (≈ 50 % of ^13^CO_2_) and for the next three weeks. ^13^C excess shows ^13^C abundance in the labelled leaf reduced by natural abundance of ^13^C in the leaves before labelling. Thick lines represent mean and shaded area one standard deviation. N=4.

**Figure 4.**
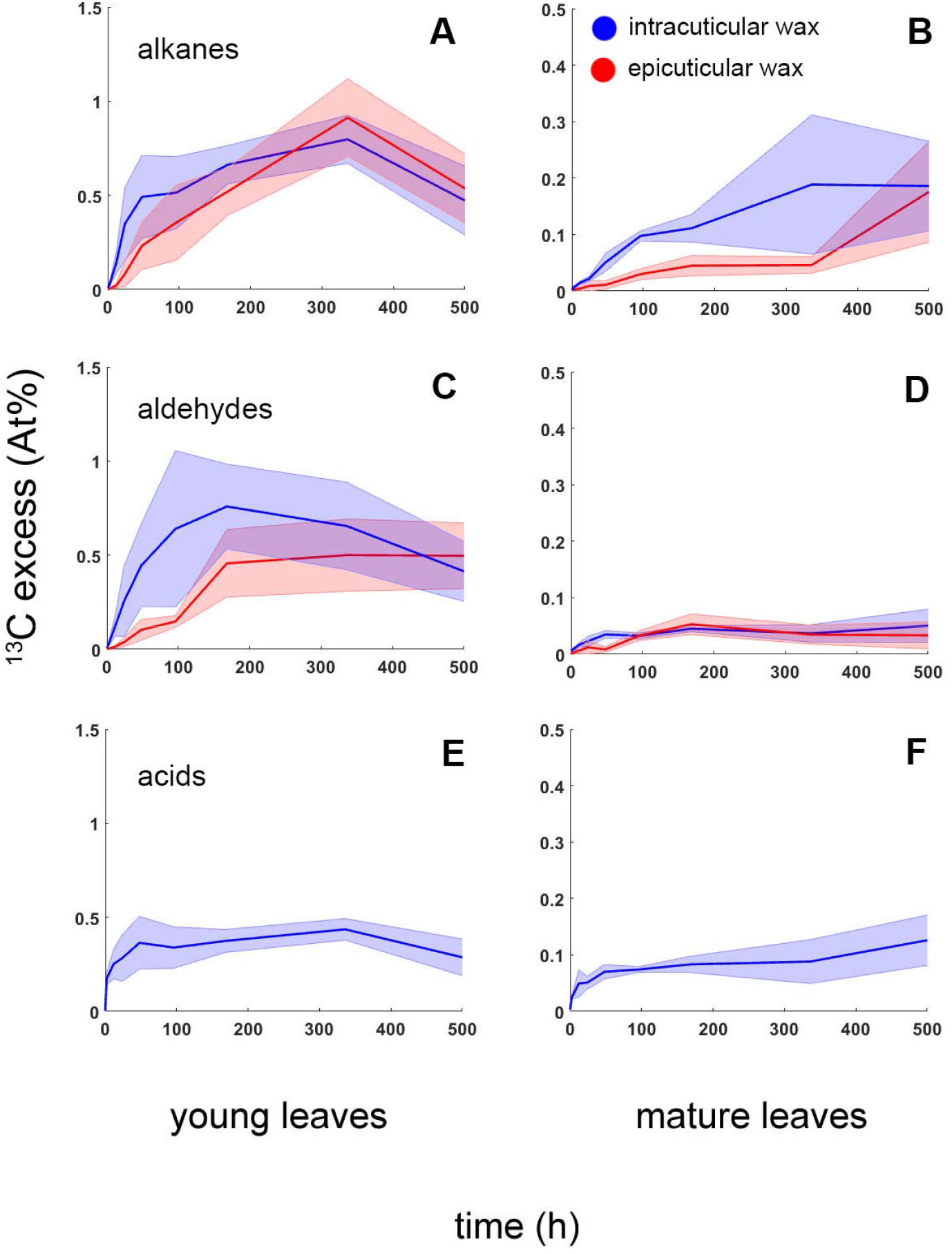
Dynamics of ^13^C abundance in main classes of wax compounds on *Clusia rosea* leaves. n-Alkanes (A, B), aldehydes (C,D) and fatty acids (E,F) of intracuticular (blue) and epicuticular (red) leaf wax are shown for young (A,C,E) and mature (B,D,F) leaves before labelling, after 2 h of CO_2_ assimilation in air highly enriched in ^13^CO_2_ (≈ 50 % of ^13^CO_2_) and for the next three weeks. Thick lines represent mean and shaded area one standard error of mean. N=4. Amount-weighted ^13^C enrichment values of particular compounds were averaged. C29, C31, C33 (and much less C27, C28, C30, C32, C35, C37) alkanes, C28, C30, C32, C34 and C36 aldehydes and C16 and C18 fatty acids (only in IW) were detected and quantified. Note different y-axis scaling for young and mature leaves.

### Compound-specific ^13^C dynamics in waxes and cutin matrix

Alkanes and aldehydes showed roughly similar ^13^C dynamics, consistent with a common synthetic pathway, the ‘alkane pathway’ (Bernard & Joubès, 2013). They were more strongly and more rapidly labelled in young leaves than in mature ones (Fig. 4). The fatty acids (C16 and C18) were labelled rapidly in the first hours after ^13^CO _2_ treatment and were found only in IW, which is very likely due to the non-cuticular origin of these compounds indicated by the extremely tight correlation of ^13^C excess between leaf sides (see Fig. S8).

IW alkanes, particularly in mature leaves, exhibited much faster ^13^C enrichment in the early period after labelling compared with EW alkanes, suggesting deposition in the matrix before the cuticle surface. Absolute ^13^C enrichment of a compound is a function of (i) the rates of synthesis, transport and deposition (ii) the size of the unlabelled pool already present, and (iii) the rate of wax loss (e.g. by abrasion) and reabsorption backward into epidermal cells. We assume in this study that (iii) was negligible during the three-week duration of the experiment. ^13^C enrichment recalculated according equation (1) as new wax deposition (NWD) shows that alkanes were preferentially allocated in IW in young leaves, whereas in mature leaves the NWD was equal for IW and EW (Fig. 5). This recalculation also demonstrated about the same NWD of alkanes for young and mature leaves (Fig. 5, Fig. 8E,F). The NWD of aldehydes was negligible due to their low abundance. We could not find any time-dependent pattern in either ^13^C abundance or NWD between the differently elongated chains (e.g. C29, C31, and C33 alkanes) and between alkanes and aldehydes (data not shown).

**Figure 5.**
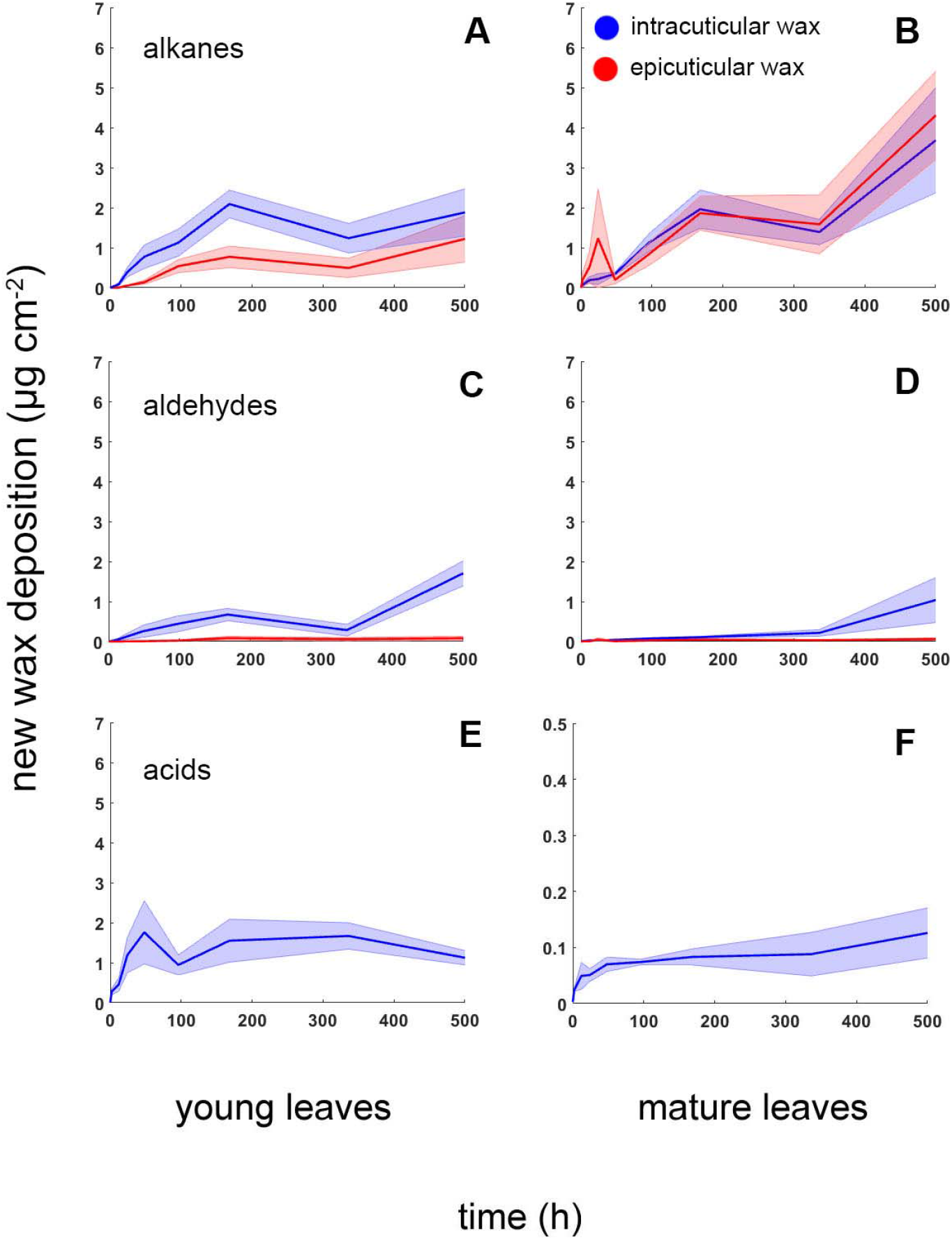
New wax deposition (NWD) in main compound classes (alkanes - A, B; aldehydes - C, D and fatty acids - E, F) of *Clusia rosea* intracuticular (blue) and epicuticular (red) leaf wax for young (A,C,E) and mature (B,D,F) leaves before labelling, after 2 h of CO_2_ assimilation in air highly enriched in ^13^CO_2_ (≈ 50 % of ^13^CO_2_) and for the next three weeks. NWD was calculated as a product of ^13^C enrichment of the particular compound class (values in Fig. 4), amount of this compound class (Table 1) divided by ^13^C enrichment in soluble leaf fraction (Fig. 2), see Material and methods. Thick lines represent mean and shaded area one standard error of mean. N=4.

The cutin matrix (MX) contained significant and highly variable amounts of new carbon ^13^C (up to 1.2 At%) only in young leaves (Fig. 3, Fig S8). Moreover, the traces of ^13^C excess in mature leaves (< 0.1 At% in most cases) may be the remnants of incompletely removed or even matrix-bound wax (Hauke & Schreiber, 1998).

### Differences between the leaf sides

Despite the different adaxial (astomatous) vs. abaxial (stomatous) leaf/cuticle morphology (Fig. 1), MX area density and EW and IW loads were similar for both leaf sides (Table 1). The chemical composition of EW and IW differed slightly but consistently between leaf sides. Both EW and IW compounds were significantly shifted towards longer carbon chains on the abaxial side (see greater average carbon length - ACL parameter in Table 1). The ratio of alkanes C29 and C33 is a major determinant of ACL in *C. rosea* leaves (Fig. S7). These ratios for adaxial and abaxial surfaces form distinct 2D data clusters (with a lower proportion of C29 in the abaxial cuticles), particularly in mature leaves (Fig. 6A,B). By contrast, the new carbon deposition expressed as ^13^C excess in both alkanes showed no preference between adaxial and abaxial sides or IW and EW (Fig. 6C,D). Thus, the new wax deposition (NWD) is proportional to the amount of compound, i.e. higher for C29 alkane on the adaxial surface and higher for C33 alkane on the abaxial surface (Fig. 6E,F).

**Figure 6.**
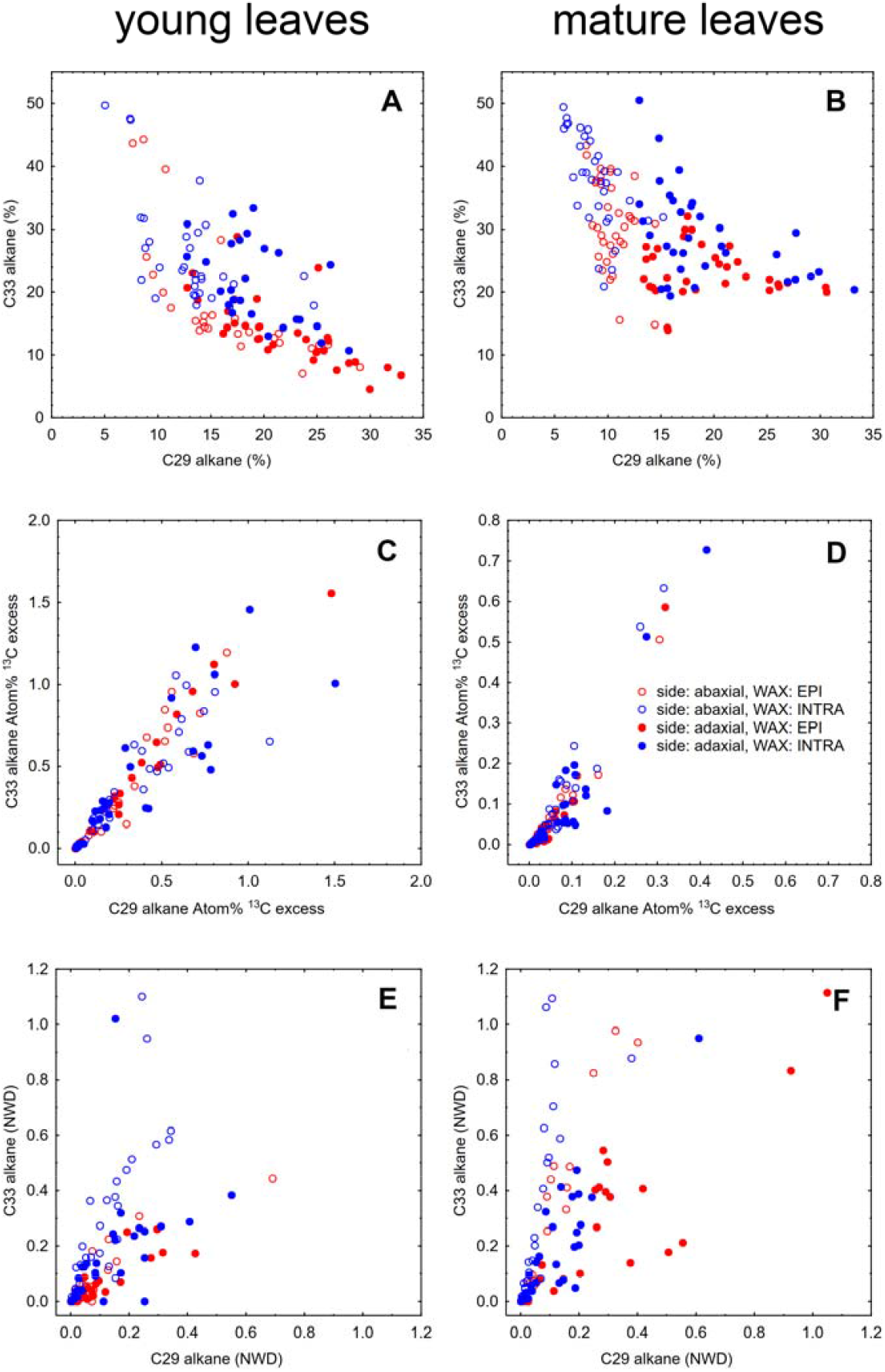
Differences between adaxial and abaxial and intracuticular and epicuticular waxes. Percentage of C29 alkane (nonacosane, x-axis) and C33 alkane (tritriacontane, y-axis) of all alkane wax content (A, B), ^13^C excess (C, D) and new wax deposition, NWD (μg cm-^2^, E, F) in those compounds for young (left column) and mature (right column) leaves. Waxes extracted from abaxial (lower) leaf side are shown as red symbols, waxes from adaxial (upper) leaf side as blue symbols. Open symbols represent epicuticular waxes, closed symbols intracuticular waxes, respectively. N=72.

### Effect of collodion treatment on leaf gas exchange

The side-specific gas exchange measurements were carried out before and after leaf treatment with collodion with the aim to assess the role of EW in gas exchange and any effect of collodion on stomata. CO_2_ assimilation (Fig. S5A,B) and stomatal conductance (Fig. S5C,D) measured at the abaxial (stomatous) side of the leaf were highly variable (especially in mature leaves). However, the collodion treatment had little, if any, effect on photosynthesis and stomatal behavior. Thus, the collodion treatment had no deleterious effect on the leaves and stomata of *C. rosea*. On the other hand, we could not detect any gas exchange, of either CO_2_ or H_2_O, on the adaxial (astomatous) side of the leaves, regardless of whether or not the collodion treatment had been applied (data not shown). Thus, the removal of EW by collodion did not significantly increase cuticular permeability.

## Discussion

This is, to the best of our knowledge, the first study of the dynamics of all major parts of the plant cuticle and of individual chemical compounds using a ^13^C tracer. We have shown that photosynthesis in air highly enriched in ^13^CO _2_ resulted in a substantial labelling of the active metabolite pool and subsequently of cutin and wax in autograph tree (*C. rosea*) and broccoli (*Brassica oleracea* var. *gemmifera*).

### Collodion is able to discriminate between EW and IW and does not harm leaf gas exchange

There is extensive discussion in the literature about the best method to sample EW and leave IW as intact as possible. Collodion solution (Haas & Rentschler, 1984), gum Arabic (Jetter & Schäffer, 2001b) and frozen water/liquids (Ensikat *et al*., 2000) are among the most common techniques. Individual research groups mostly prefer the one they have developed. Collodion is a solution of ca. 4 % nitrocellulose in 1:1 diethyl ether and ethanol. Some researchers point out that these organic solvents may mobilize part of IW and recommend an aqueous solution of gum Arabic instead (Jetter *et al*., 2000). Since gum Arabic is harder to work with, contains organic contaminants and is not suitable for working on isolated cuticles (Zeisler & Schreiber, 2016), collodion appears to be a more versatile option. Here we confirm that EW and IW may be distinguished in *C. rosea* leaves by collodion sampling. Moreover, only two consecutive strips were sufficient to sample more than 90 % of the EW (Fig. S3). At the same time, no gas exchange of the upper (astomatous) side of leaves before and after the collodion treatment was detectable (with system noise: ≈0.001 mol m-_2_ s^−1^ for stomatal conductance and ≈0.1 μmol m-_2_ s^−1^ for CO_2_ exchange). Thus, we confirm that collodion does not have a detrimental effect on living leaves of *C. rosea* and their stomata (Fig. S5). This is in agreement with previous experiments on *Prunus laurocerasus* leaves and cuticles, where collodion-derived EW contained only trace amounts of triterpenoids found exclusively in IW and up to five consecutive treatments did not disrupt the cuticle and its conductance (Zeisler & Schreiber, 2016).

### Wax is deposited inside the cuticle before being exported to the leaf surface - the leaf lifespan perspective

New (^13^C) carbon was detectable first (as early as 2h after labelling) in IW, which remained enriched in comparison to EW for most of the experimental duration (Fig. 3), particularly in alkanes (Fig. 4). Two weeks after labelling, ^13^C excess in EW surpassed that of IW in young leaves. In mature leaves, ^13^C changes were smaller and slower (Fig. 3, Fig. 4) due to the high wax load (Table 1). The ^13^C excess more accurately depicts the amount of deposited wax in the two pools examined when converted to NWD (Fig. 5). This implies that the MX must first be filled with wax and only small amount is exported to the leaf surface in young leaves (Fig. 5A). The very low EW load of young leaves further supports this conclusion. Surprisingly, mature (last year) leaves with approximately ten times higher EW coverage (Table 1) allocated approximately equal amounts of alkanes in IW and EW (Fig. 5B). Recently, IW, not EW, has been identified as the main component of the gas transport barrier of leaf cuticles (Zeisler & Schreiber, 2016; Zeisler-Diehl *et al*., 2017; Zeisler-Diehl & Müller, 2018). Sealing the young leaf against desiccation stress by IW seems to be the primary effort, followed by export of additional wax (EW) to the cuticle surface, which serves other cuticle-specific purposes. Mature leaves, on the other hand, may have saturated MX with IW during the earlier stages of their development, resulting in the increased deposition of EW providing an “armor” against pathogens, xenobiotics and/or UV radiation penetrating the leaf (Lewandowska *et al*., 2020). Interestingly, the rate of wax synthesis appears to be roughly comparable in young and mature leaves (Fig. 5). Thus, cuticular waxes are renewed in *C. rosea* throughout leaf ontogeny (except perhaps during senescence) despite the rate of loss of waxes from the leaf surface, and thus the exact rate of wax turnover, could not be inferred reliably from this study.

### Wax renewal as a strategic trait reflecting climatic factors and plant/leaf life strategies

Lipid synthesis is one of the most energy-demanding processes in living cells (Buchanan *et al*., 2015). The cuticle is usually thin but its mass is not negligible. Consider a “typical” mesic leaf with a thickness of 0.2 mm. Two cuticles, on both sides, each 1 μm thick, represent 1 % of the leaf cross-section. In addition, the weight of a leaf is usually reduced by 90 % or more after drying, while the weight of the cuticle, containing mainly lipids, remains virtually unchanged. The cuticle therefore represents a significant proportion of leaf dry matter (up to 24 %; on average 5 to 10 %; Onoda *et al*., 2012). Selective pressure should optimize cuticle dynamics in response to the environment and leaf longevity. It may explain the abovementioned controversy in published data on wax renewal during leaf expansion or throughout the leaf lifespan (Sachse *et al*., 2009; Kahmen *et al*., 2011; Gao *et al*., 2012). Four distinct “functional groups” with respect to EW regeneration were also found by Neinhuis *et al*. (2001) using SEM; however, ecological implications were not identified. Our recent data suggest that young *Brassica oleracea* leaves allocate much more wax per leaf area (by at least one order of magnitude) than leaves whose expansion has ceased (Fig. S9). Thus, annual leaves may save expending high-energy lipid resources on the renewal of wax in comparison to leaves with higher longevity (e.g., *C. rosea*), which must continuously allocate part of their energy and metabolites to wax turnover.

### Wax elongation is very fast, and fatty acids detected are not of cuticular origin

The predominant compounds in the wax of *C. rosea* are various long-chain aliphatic compounds C25 to C38 in length, mainly C29, C31 and C33 n-alkanes (nonacosane, hentriacontane and tritriacontane). Our efforts to detect differences in the relative amounts of newly synthesized compounds with different chain lengths, or between alkanes and aldehydes, did not provide a clear picture. There was a slight trend indicating that the ^13^C excess is higher for the shorter chain IW two hours after labelling (EW was not sufficiently labelled at this time, data not shown). Thus, the elongation of fatty acids and formation of compounds with various functional groups and their transport to the uppermost leaf surface was surprisingly rapid. Shedding more light on these issues will require additional research.

Fatty acids, mainly C22 and longer, are frequently found in plant cuticles (Haas & Rentschler, 1984; Hauke & Schreiber, 1998; Bernard & Joubès, 2013). On the other hand, effective adsorption of C16 and C18 fatty acids from tissue slurries was observed already in 1986 during enzymatic cuticle isolation (Schönherr & Riederer, 1986). We found substantial amounts of hexadecanoic and octadecanoic acids in IW of isolated cuticles but not in in EW (Fig. 2). Based on ^13^C data, we argue these acids were not of cuticular origin. The extremely close correlation between adaxial and abaxial ^13^C excess observed (Fig. S8, r^2^ > 0.99) is possible only if the following two conditions are met: 1/ the pool of fatty acids is the same, thus originating from leaf tissue, and 2/ the precision of the isotopic measurement is high. This result proves the accuracy of our method and demonstrates that short fatty acids are not a major constituent of *C. rosea* cuticles.

### Cuticle biosynthesis is tightly coordinated between leaf sides

The abundances of major compounds and ^13^C dynamics in the cuticles on opposite sides of *C. rosea* leaves were tightly correlated (Fig. 6, Fig. S8), which contrasts with the differing morphology (Fig. 1) and presumably contrasting environmental conditions on opposite sides of the leaves (Bird & Gray, 2003a). Moreover, *C. rosea* leaves are hypostomatous, implying a gradient of photosynthesis and CO_2_ across the leaf (Terashima *et al*., 2011). This side-specific coordination of cuticle development and renewal, first reported here using ^13^C tracer, may be anticipated but mechanisms are unknown.

Probably even more puzzling are the subtle and consistent side-specific differences in abundance of some compounds. For example, the ratio of C29 and C33 alkanes can predict the leaf side of *C. rosea* with nearly 100% confidence, particularly in mature leaves (Fig. 6B). The side-specific ^13^C excess, in turn, was very similar for both leaf sides (Fig. 6C,D). This implies a proportionality of the rate of synthesis to the amount of a particular compound (Fig 6E,F) and likely maintains the leaf-side specific C29/C33 ratio during development. Recently, we have found an even more striking contrast in EW of pepper. The ratio of the abundances of C29 to C33 alkanes was 0.98 ± 0.2 for the adaxial side but only 0.28 ± 0.20 for the adaxial side (N=60, Fig. S10). These side-specific differences in cuticular wax chemistry have been known for a long time (Holloway *et al*., 1977) but the physiological functions and mechanisms maintaining these contrasts are not clear. One possible explanation could arise from the side-specific presence or frequency of stomata. In most plant species, stomata are located preferentially on the abaxial leaf side, and the cuticle covering guard cells differs from the pavement cell cuticle in thickness and wax composition (Karabourniotis *et al*., 2001; Bird & Gray, 2003b). If the fraction of the leaf area taken up by guard cells increases, we might expect that the discrepancy in wax composition between opposite leaf sides will rise. However, this hypothesis remains to be tested in future studies. To the best of our knowledge, to date, only biotic interactions - microbial preferences for a particular leaf side - have been correlated with side-specific cuticular wax chemistry (Gniwotta *et al*., 2005).

In conclusion, ^13^C isotope labelling and tracing let us show that (i) both epicuticular and intracuticular waxes are renewed during the whole leaf lifespan; (ii) wax compounds are deposited in the cuticle as intracuticular wax (IW) first and may be exported to the leaf surface as epicuticular wax (EW) later on; (iii) only young, still expanding leaves incorporate new carbon into the cutin matrix (MX); and (iv) differences in wax amount and composition between leaf sides are typically small but consistent, suggesting coordinated adaxial and abaxial cuticle formation. Methodically, we have shown that collodion is able to effectively discriminate between EW and IW and does not adversely affect leaves and stomata, at least in *Clusia rosea*, thus making it suitable for in-vivo wax regeneration studies.

## Acknowledgement

We express our thanks to Petra Fialová for technical assistance with plant cultivation and Ladislav Marek for bulk-isotope analyses.

This research was supported by Czech Science Foundation grant No. 18-14704S.

## Author contributions

JŠ and JK planned and designed the research (gas exchange, ^13^C labelling and measurement), JK processed data and wrote the manuscript; TK sampled leaves and prepared samples for isotope analyses; JJ planned and designed the sampling during the experiment, planned and co-ordinated the pilot measurements, sample harvesting and preparing for isotopic analyses; BA sampled EW with collodion, isolated cuticles, obtained IW and processed some data, JB identified wax compounds using GC-MS-TOF. All authors approved the manuscript.

The following Supporting Information is available for this article:

**Fig. S1 ^13^CO_2_ labelling facility.**

**Fig. S1 ^13^CO_2_ labelling facility.**

**Fig. S2 Spectral output of LED modules used for plant acclimation and to drive photosynthesis during** ^**13**^**CO**_**2**_ **labelling**.

**Fig. S3 Relative amounts of different epicuticular wax (EW) compounds removed by stripping with three subsequent collodion layers applied to the leaf surface (≈ 30 cm**_**2**_**) of *Clusia rosea* plants (adaxial and abaxial sides pooled)**.

**Fig. S4 Two Li-6400XT set-up**.

**Fig. S5 Photosynthetic induction of *Clusia rosea* leaves subjected to collodion treatment. Abaxial gas exchange**.

**Fig. S6 Statistical distribution of cuticular waxes (upper four panels) and epicuticular wax-free, enzymatically isolated cuticles (lower two panels)**.

**Fig. S7 Typical chromatograms of epicuticular waxes of abaxial and adaxial sides of *Clusia rosea* leaves**.

**Fig. S8 Correlation of** ^**13**^**C excess between adaxial and abaxial cuticle compound classes and dewaxed cutin matrix (MX)**.

**Fig. S9 Epicuticular wax of *Brassica oleracea var. gemmifera* during pilot** ^**13**^**C labelling experiment**.

**Fig. S10 Side specific epicuticular wax of *Capsicum annuum* (pepper)**.

