## Supplemental file for "Cuticular wax, but not the cutin matrix, is renewed during the lifespan of *Clusia rosea* leaves *^13^CO_2_ labelling and gas exchange study*"

The following Supporting Information is available for this article:

**Fig. S1 $^{13}CO_2$ labelling facility.**

**Fig. S2 Spectral output of LED modules used for plant acclimation and to drive photosynthesis during $^{13}CO_2$ labelling.**

**Fig. S3 Relative amounts of different epicuticular wax (EW) compounds removed by stripping with three subsequent collodion layers applied to the leaf surface (≈ 30 $cm^2$) of *Clusia rosea* plants (adaxial and abaxial sides pooled).**

**Fig. S4 Two Li-6400XT set-up.**

**Fig. S5 Photosynthetic induction of *Clusia rosea* leaves subjected to collodion treatment. Abaxial gas exchange.**

**Fig. S6 Statistical distribution of cuticular waxes (upper four panels) and epicuticular wax-free, enzymatically isolated cuticles (lower two panels).**

**Fig. S7 Typical chromatograms of epicuticular waxes of abaxial and adaxial sides of *Clusia rosea* leaves.**

**Fig. S8 Correlation of $^{13}C$ excess between adaxial and abaxial cuticle compound classes and dewaxed cutin matrix (MX).**

**Fig. S9 Epicuticular wax of *Brassica oleracea var. gemmifera* during pilot $^{13}C$ labelling experiment.**

**Fig. S10 Side specific epicuticular wax of *Capsicum annuum* (pepper).**

**Fig. S1 $^{13}CO_2$ labelling facility.** Gas-tight box (internal dimensions 60 x 60 x 60 cm and volume of 190 L) with septum-sealed inlet and electric fan inside to reduce boundary layer resistance. The box with plants inside was illuminated by linear LED sources (for spectral output see Fig. S2). For complete labelling procedure see Material and Methods.

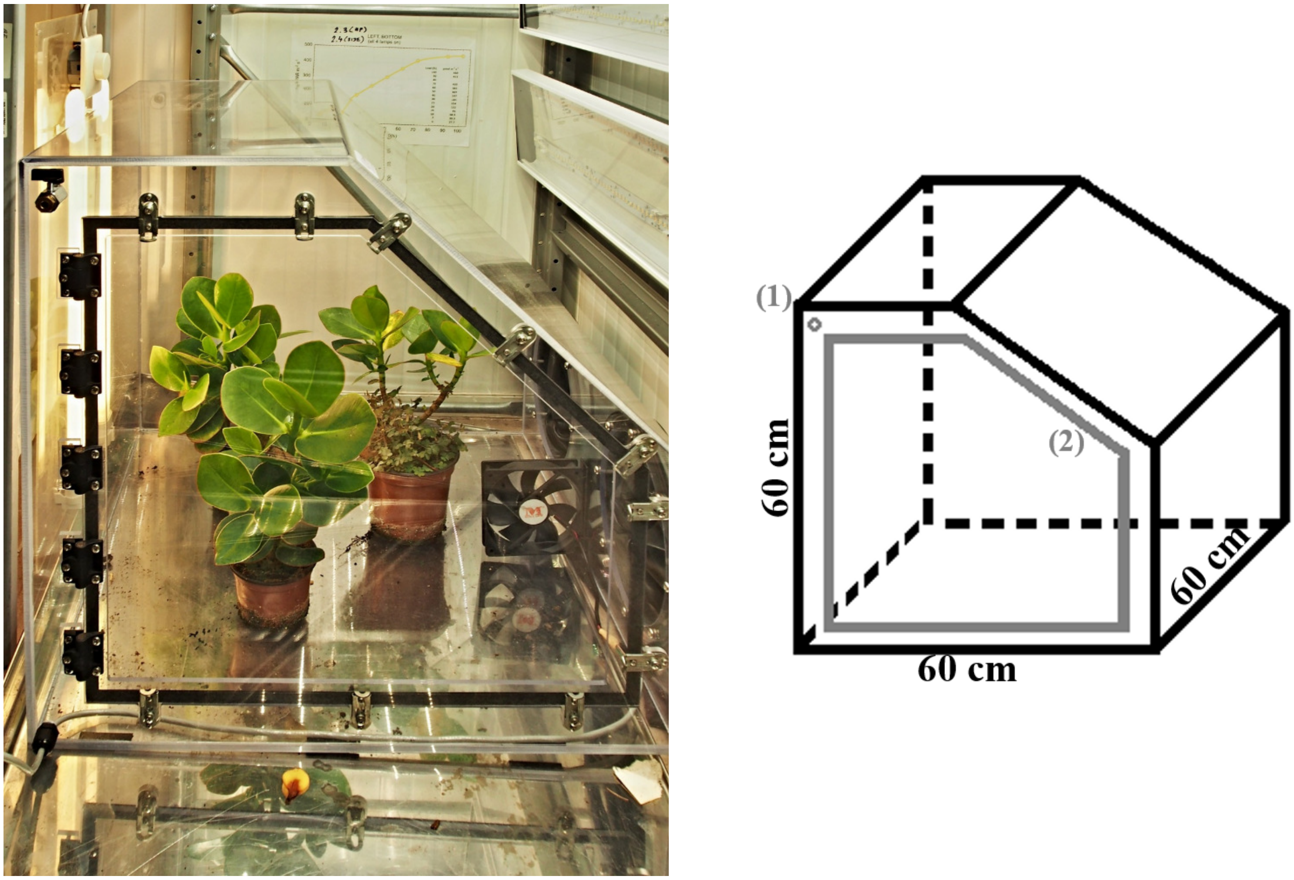

**Fig. S2 Spectral output of LED modules used for plant acclimation and to drive photosynthesis during $^{13}CO_2$ labelling.** LED chips emit blue light (c. 450 nm) and the luminophore converts part of these photons to a broad band of photons of longer wavelength. Measured with handheld spectrometer SpectraPEN mini (PSI, Drasov, Czech Republic).

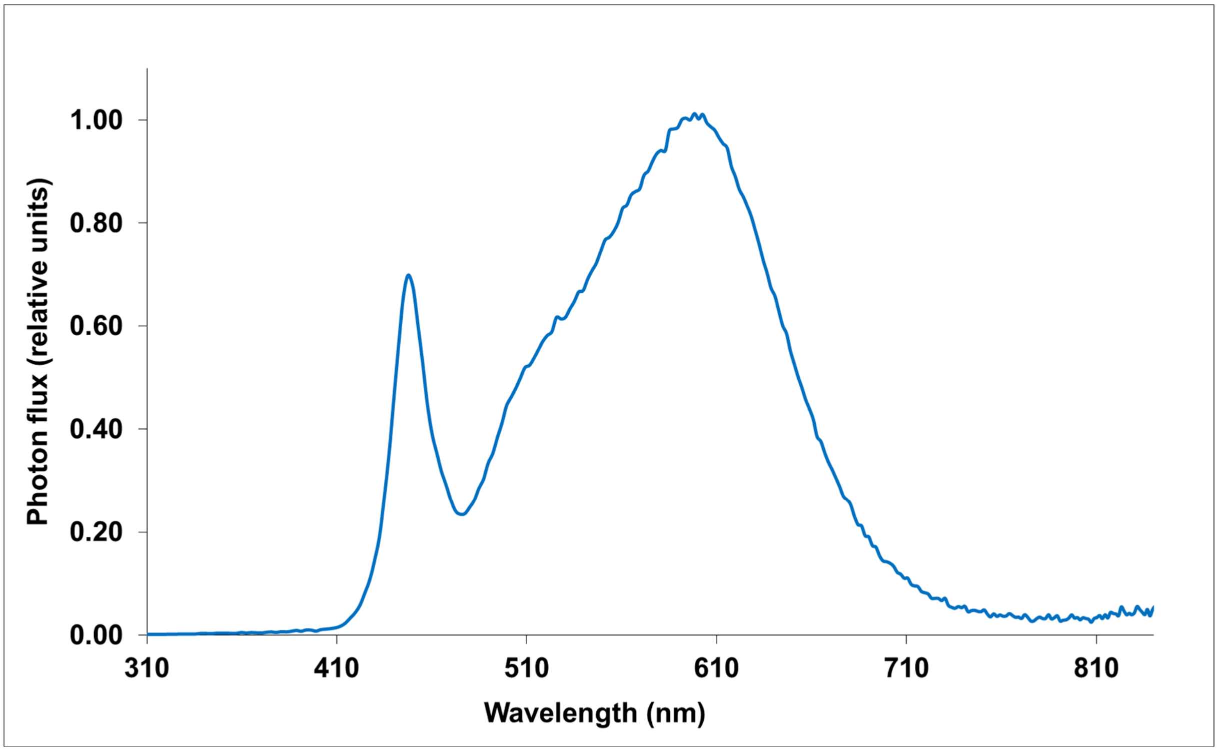

**Fig. S3 Relative amounts of different epicuticular wax (EW) compounds removed by stripping with three subsequent collodion layers applied to the leaf surface (≈ 30 cm$^2$) of *Clusia rosea* plants (adaxial and abaxial sides pooled).** Cumulative amount after third strip (L1+L2+L3) was set to 100 %. Several minor compounds reached 100 % already in second strip, which means that chromatographic peak from third-strip was under limit of quantification. Each strip was extracted in 4 ml of n-hexane on the roller overnight, then the strip was removed and the sample concentrated to 1 ml and analyzed by GC-IRMS (see Material and Methods). Alkanes are slightly more prone to be removed by collodion in comparison to aldehydes.

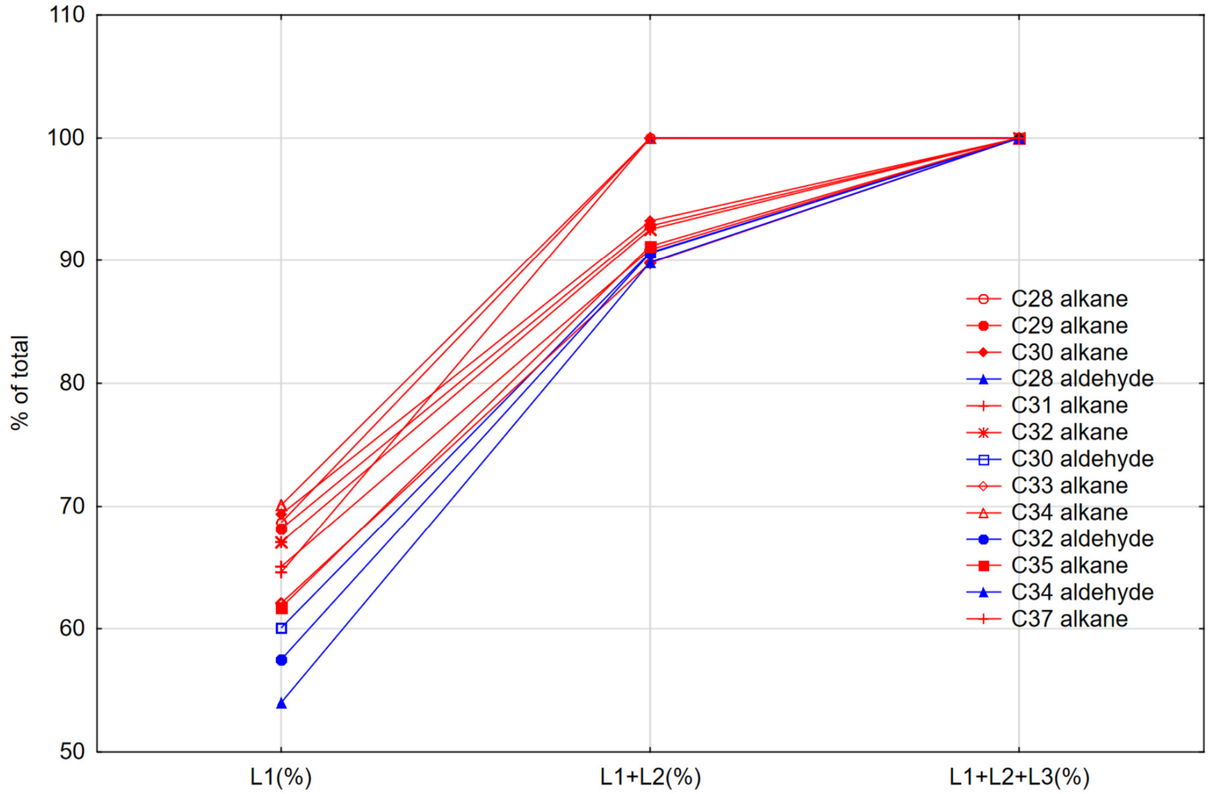

**Fig. S4 Two – Li-6400XT set-up.** Two LI-6400XT units were used and their 2x3 cm leaf chambers were arranged as a "tandem", one in an inverse position. Left unit (in this view) contained only the bottom part of one chamber and was measuring the abaxial (stomatal) leaf side. Right unit consisted of only the upper part of the other chamber with LED light source and was measuring adaxial (astomatous) side of *Clusia rosea* leaf. We originally optimized this set-up for measurement of amphistomatous leaves where not only diffusional but also bulk flow may occur across the leaf (continuum of stoma – intercellular air – stoma). Thus, we were able to monitor and maintain pressure difference between adaxial and abaxial unit of less than 0.1 mBar (10 Pa), which is important to rule out bulk flow across a leaf. On the other hand, hypostomatous leaves such as *Clusia rosea* here have a practically gas-tight adaxial cuticle and side-specific gas exchange measurements are much less demanding. Overall, we were able to measure adaxial and abaxial, and water and $CO_2$ leaf fluxes separately with high precision.

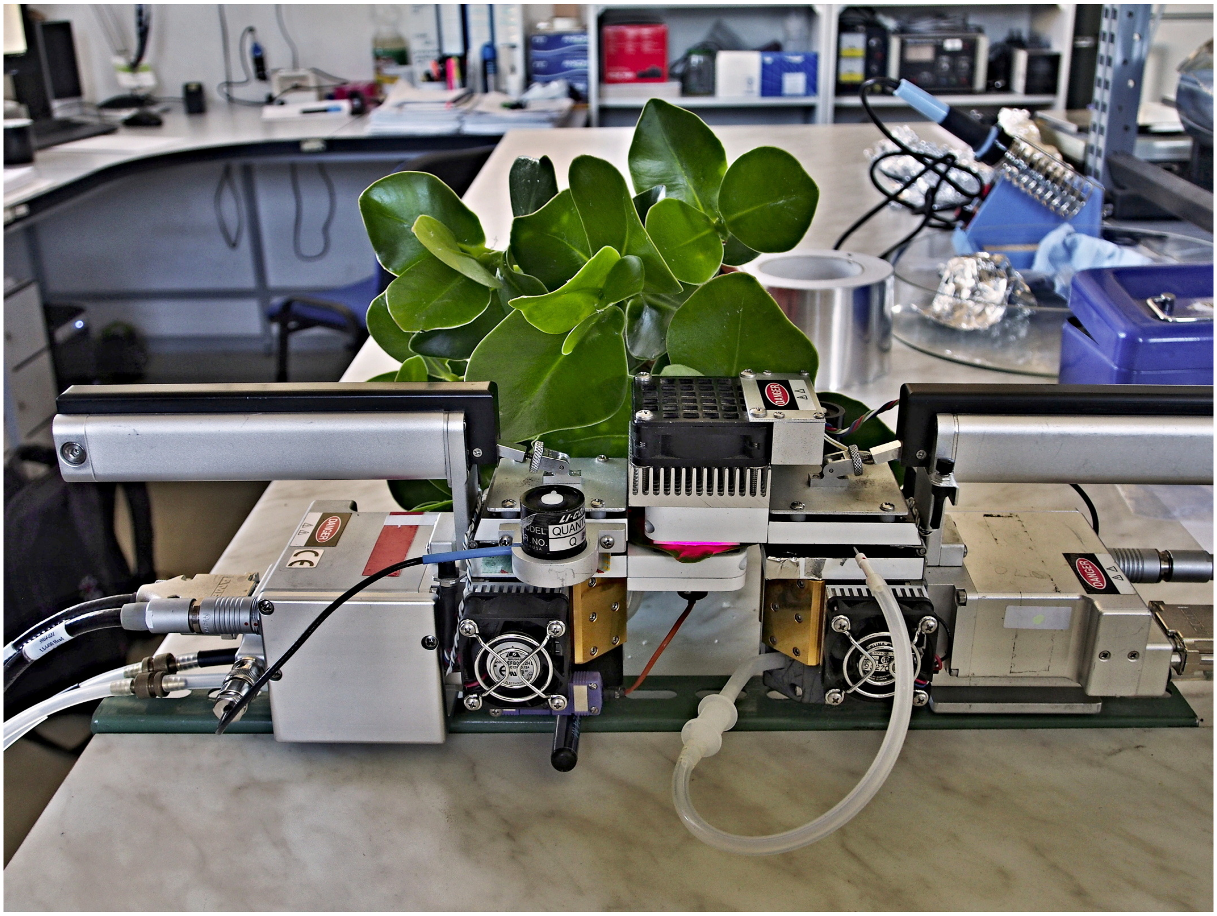

**Fig. S5 Photosynthetic induction of *Clusia rosea* leaves subjected to collodion treatment. Abaxial gas exchange.** An area of 2 x 3 cm of a leaf of a dark-acclimated plant (1 h) was inserted in the ”tandem” chamber of the two LI-6400XT set-up (see Fig S6). After several minutes in the dark (to reach system steady-state), light was set to 1000 µmol m$^{-2}$ s$^{-1}$ at time 0 and $CO_2$ assimilation and stomatal conductance logged every ten seconds for about one hour. Then, the leaf was removed from the chamber and both sides were covered with collodion solution using a fine brush. After several minutes, the collodion polymerized in a thin film and was stripped off using tweezers. The same gas exchange measurement was performed on the same leaf about 24 h later. Collodion had only minor (if any) effects on leaf gas exchange and stomatal behavior. Means (thick line) and one standard deviation (shaded area) are shown. N=3. Adaxial $CO_2$ and $H_2O$ exchange were not detectable.

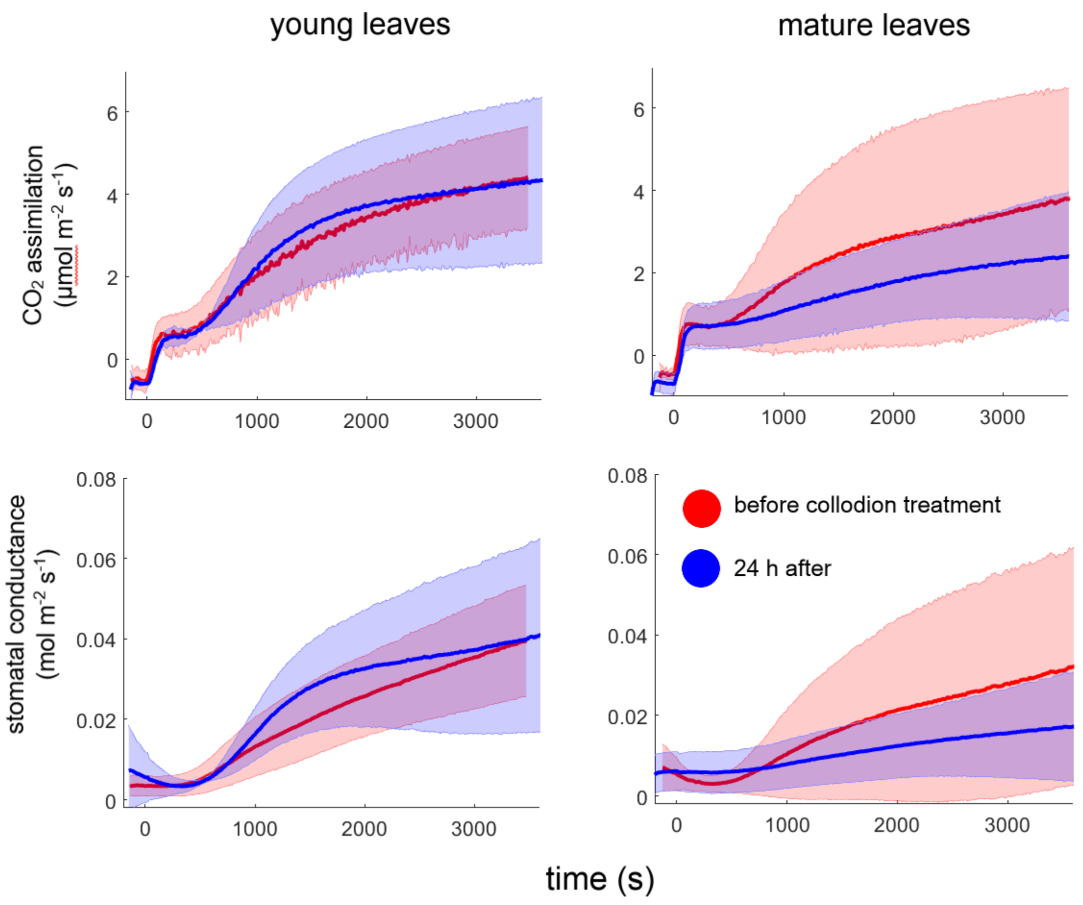

**Fig. S6 Statistical distribution of cuticular waxes (A-D) and epicuticular wax-free, enzymatically isolated cuticles (E,F).** The histograms show that 1/ epicuticular wax is very variable in total amount, preferably accumulated in mature leaves, with very low median for young leaves (the category 0 to 1 μg cm$^{-2}$ was the most common); 2/ intracuticular wax is much more uniform and distribution is almost Gaussian; 3/ area density of isolated cuticles is the most uniform variable (standard deviation less than mean).

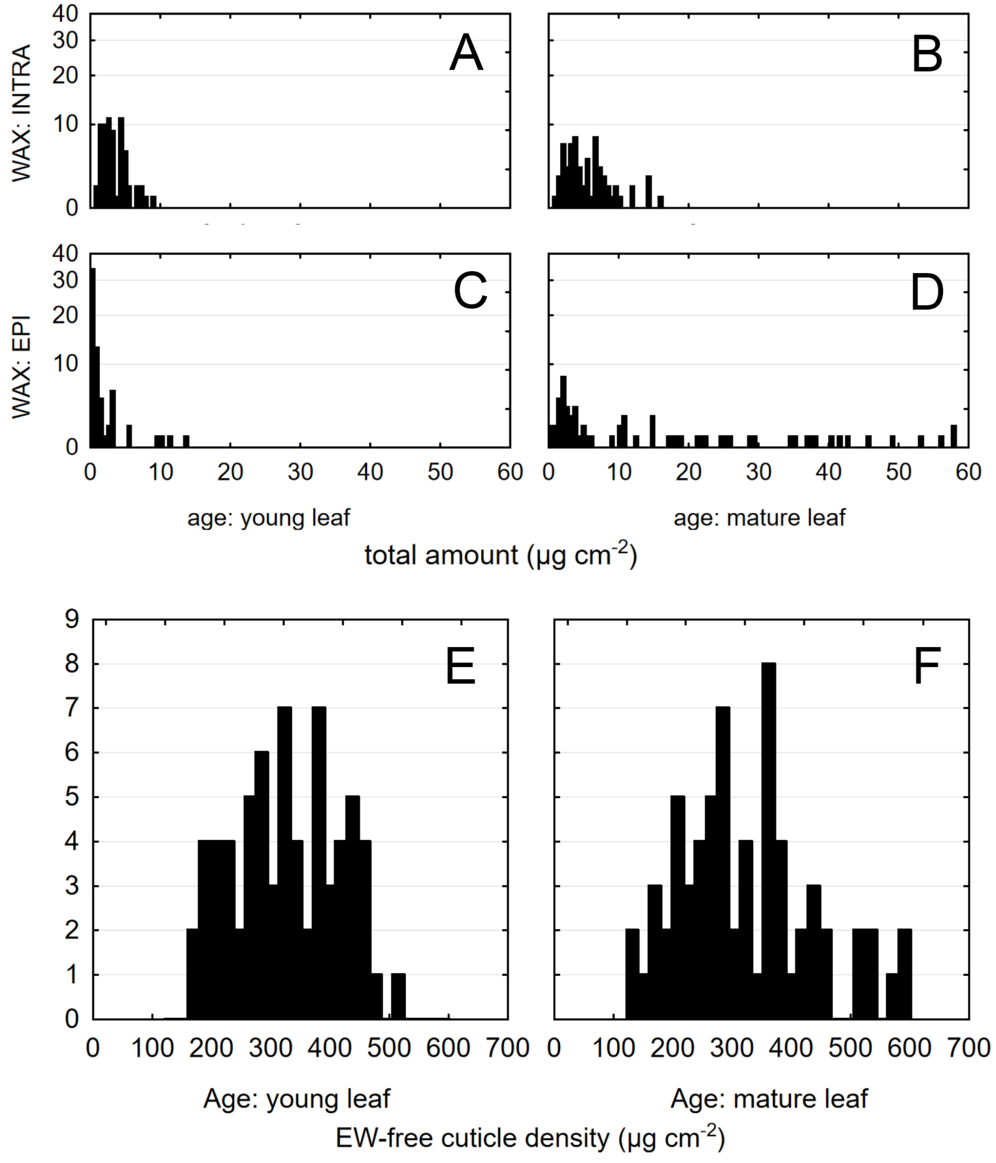

**Fig. S7 Typical chromatograms of epicuticular waxes of abaxial and adaxial sides of *Clusia rosea* leaves.** FID detector response was standardized to C31 alkane (4). Compounds are marked as follows: (1) C24 alkane (internal standard), (2) C29 alkane, (3) C30 alkane, (4) C31 alkane, (5) C32 alkane, (6) C30 aldehyde, (7) C33 alkane, (8) C34 alkane, (9) C32 aldehyde, (10) C35 alkane, (11) C34 aldehyde, (12) C37 alkane, (13) C36 aldehyde, (14) C39 alkane. The abaxial leaf side tends to accumulate longer chains, particularly C33 alkane.

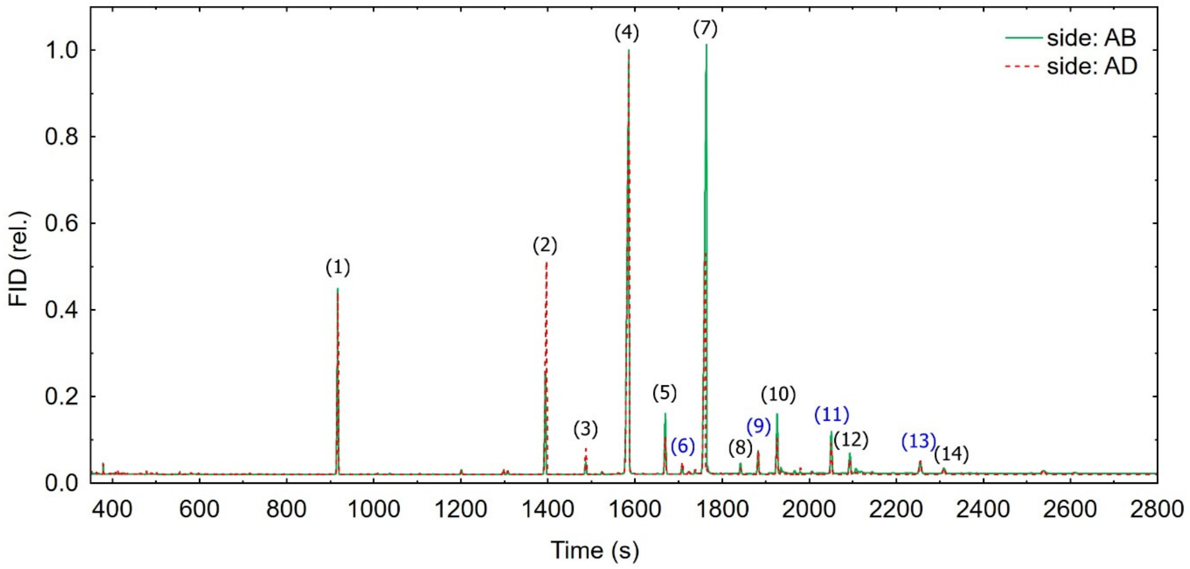

**Fig. S8 Correlation of $^{13}C$ excess between adaxial and abaxial cuticle compound classes and dewaxed cutin matrix (MX).** Coefficients of determination for each group are shown, N=72.

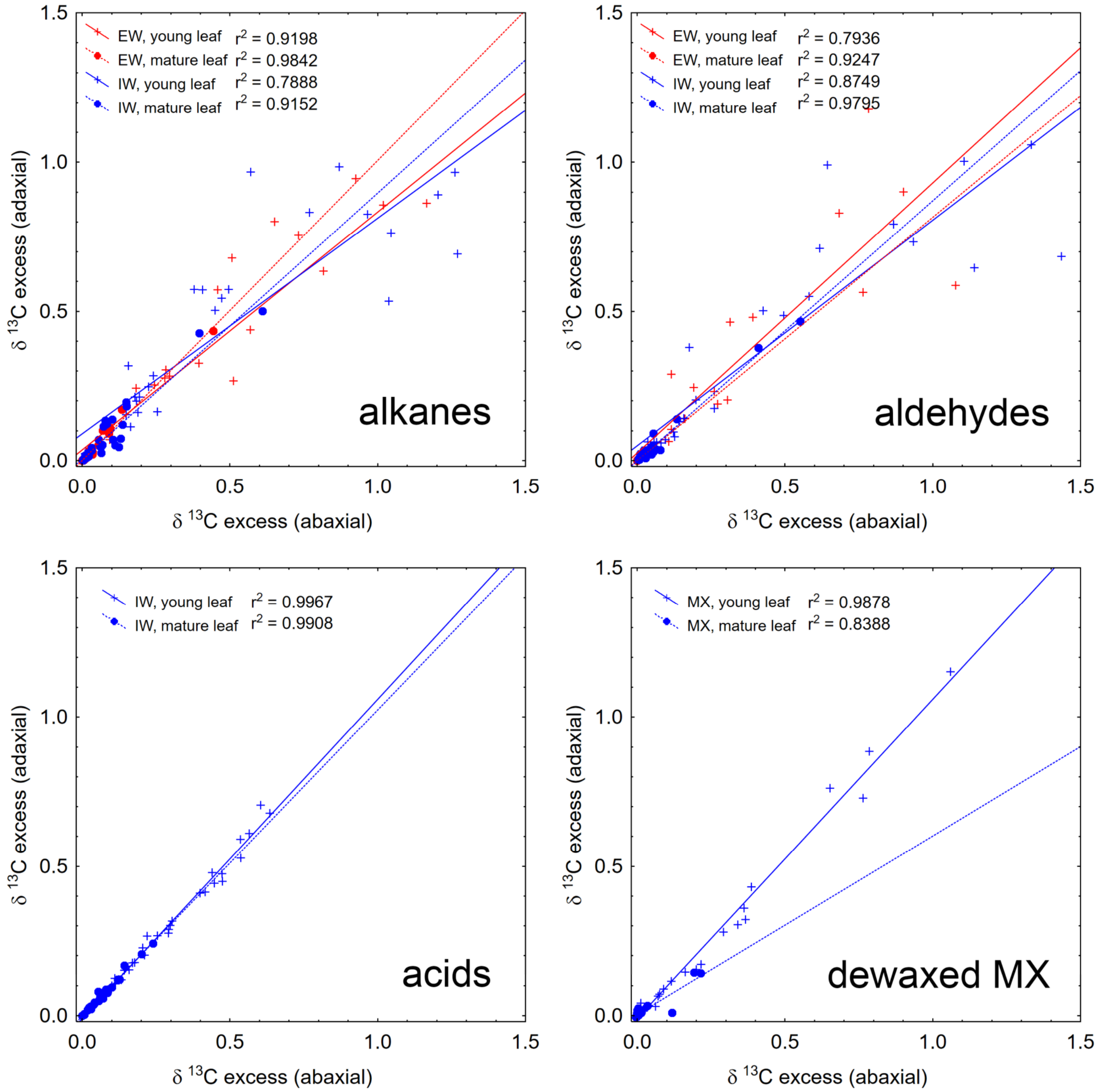

**Fig. S9 Epicuticular wax of *Brassica oleracea var. gemmifera* during pilot $^{13}C$ labelling experiment.** Labelling, sampling and sample processing protocols were the same as in the main *Clusia* experiment. Only epicuticular wax (EW) was sampled and the three most abundant compounds are shown here. Young (Y) and Mature (M) leaves did not differ in either total EW load or those of the most abundant compounds (upper panel). On the other hand, $^{13}C$ content (middle panel) and new wax deposition (bottom panel) were much higher in young leaves for all times after labelling. A slight increase at the latest times (192 and 360 h) may be due to the M leaf sampled at these times being possibly still expanding somewhat at the time of labelling. Natural $^{13}C$ content of EW was 1.07 ± 0.005 At%, N=2.

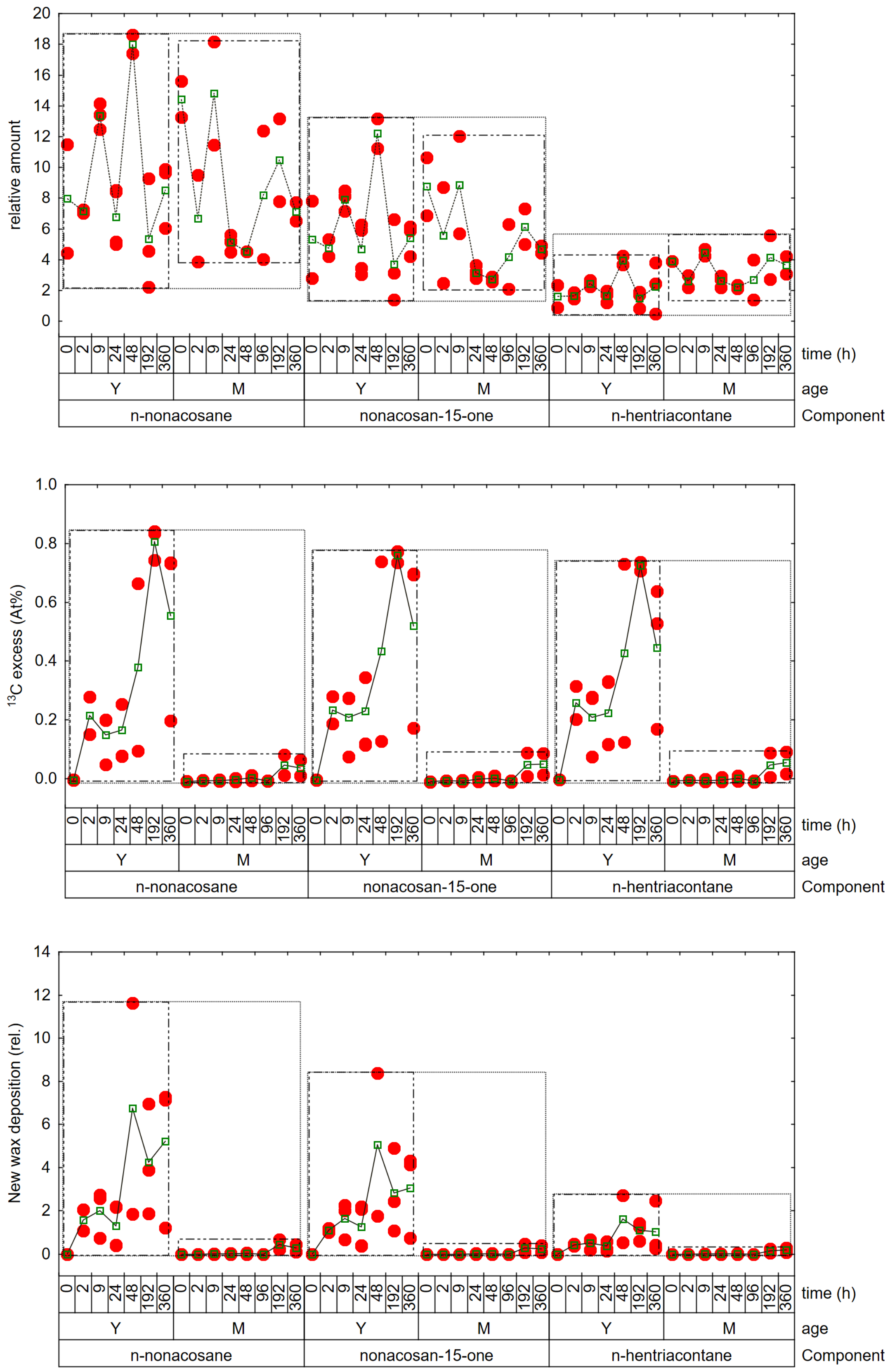

relative amount
$^{13}$C excess (At%)
New wax deposition (rel.)
time (h)
age
Component
0 2 9 24 48 192 360 0 2 9 24 48 96 192 360
Y M
n-nonacosane
nonacosan-15-one
n-hentriacontane

**Fig. S10 Side specific epicuticular wax of *Capsicum annuum* (pepper).** Sampling and sample processing protocols were the same as in the main *Clusia* experiment. Only epicuticular wax (EW) was sampled and the two abundant alkanes (C29 and C33) relative abundance are shown here.

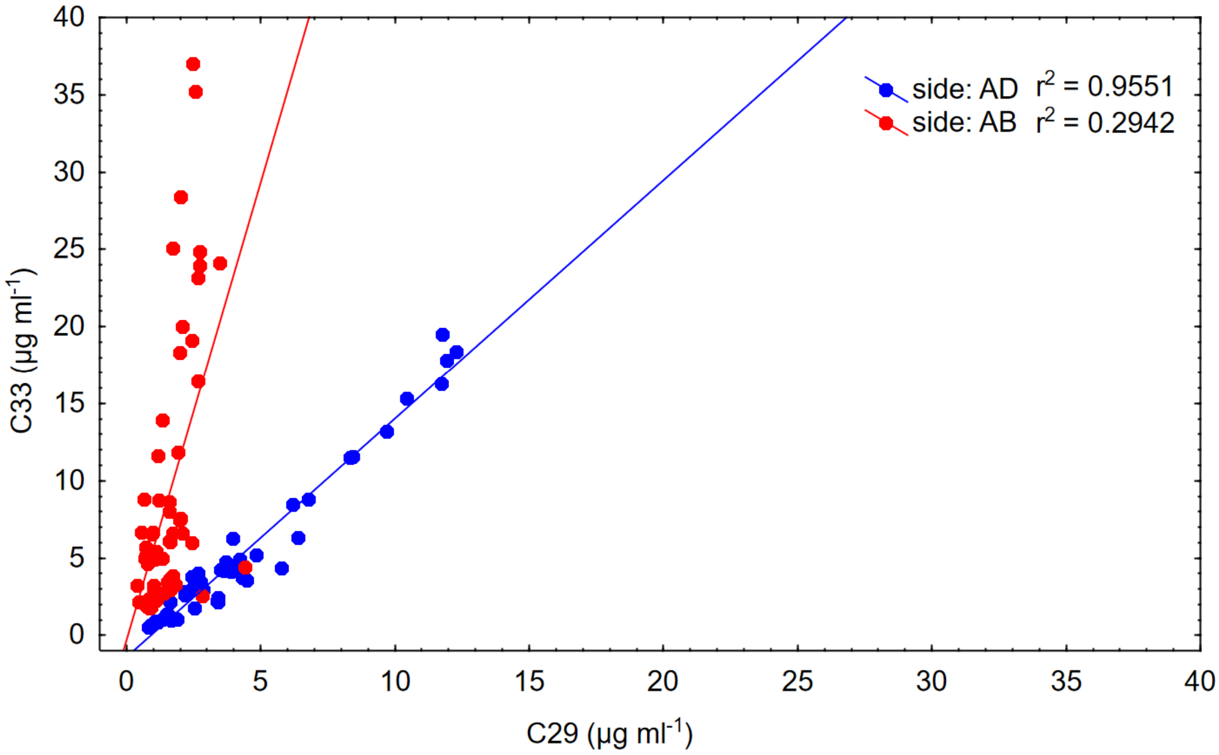
